# Neuroanatomical, electrophysiological, and morphological characterization of melanin-concentrating hormone cells coexpressing cocaine- and amphetamine-regulated transcript

**DOI:** 10.1101/2023.09.25.559204

**Authors:** Persephone A Miller, Jesukhogie G Williams-Ikhenoba, Aditi S Sankhe, Brendan H Hoffe, Melissa J Chee

## Abstract

Melanin-concentrating hormone (MCH) cells in the hypothalamus regulate fundamental physiological functions like energy balance, sleep, and reproduction. This diversity may be ascribed to the neurochemical heterogeneity among MCH cells. One prominent subpopulation of MCH cells coexpresses cocaine- and amphetamine-regulated transcript (CART), and as MCH and CART can have opposing actions, MCH/CART+ and MCH/CART− cells may differentially modulate behavioural outcomes. However, it is not known if there are differences in cellular properties underlying their functional differences, thus we compared the neuroanatomical, electrophysiological, and morphological properties of MCH cells in male and female *Mch-cre;L10-Egfp* reporter mice. Half of MCH cells expressed CART and were most prominent in the medial hypothalamus. Whole-cell patch-clamp recordings revealed differences in their passive and active membrane properties in a sex-dependent manner. Female MCH/CART+ cells had lower input resistances, but male cells largely differed in their firing properties. All MCH cells increased firing when stimulated, but their firing frequency decreases with sustained stimulation. MCH/CART+ cells showed stronger spike rate adaptation than MCH/CART− cells. The kinetics of excitatory events at MCH cells also differed by cell type, as the rising rate of excitatory events was slower at MCH/CART+ cells. By reconstructing the dendritic arborization of our recorded cells, we found no sex differences, but male MCH/CART+ cells had less dendritic length and fewer branch points. Overall, distinctions in topographical division and cellular properties between MCH cells add to their heterogeneity and help elucidate their response to stimuli or effect on modulating their respective neural networks.

## Key points

1. Topographical division between MCH/CART+ cells medially and MCH/CART− cells laterally was evident in male and female mice.
2. Female MCH cells differed in their passive membrane properties while male MCH cells differed in their capacity for excitation.
3. Male MCH/CART+ cells had less complex dendritic arborizations and tend to have slower, smaller excitatory input.

### INTRODUCTION

Melanin-concentrating hormone (MCH) is an evolutionarily conserved neuropeptide produced in the lateral hypothalamic area (LHA) of all vertebrate species (Croizier et al., 2012), though it may also be produced in extrahypothalamic regions like the lateral septum and medial preoptic area during lactation (Rondini et al., 2010; Benedetto et al., 2014; Beekly et al., 2020). MCH cells have widespread projections throughout the brain (Bittencourt, 1992), thus they have been implicated in diverse behaviours, including feeding (Qu et al., 1996; Rossi et al., 1997; Shimada et al., 1998), sleep (Verret et al., 2003; Monti et al., 2013), learning and memory (Monzon et al.,1999; Adamantidis and de Lecea, 2009; Concetti and Burdakov, 2021), mood functions like anxiety and depression (Borowsky et al., 2002; Georgescu et al., 2005; Sankhe et al., 2022), and reproduction (Gonzalez et al., 1997; Williamson-Hughes et al., 2005; Wu et al., 2009). The MCH system is also heterogeneous between males and females. For example, while MCH administration is widely recognized to increase feeding in males (Qu et al., 1996; Rossi et al., 1997; Tritos et al., 1998; Della-Zuana et al., 2002; Clegg et al., 2003; Gomori et al., 2003; Glick et al., 2009), it has little effect in females (Mogi et al., 2005; Messina et al., 2006; Santollo and Eckel, 2007; Terrill et al., 2020). The orexigenic effects of MCH are also estrogen-sensitive, as chronic, but not acute (Tritos et al., 2004), estrogen treatment blocked fasting-induced increases in *Pmch* gene expression within the LHA (Murray et al., 2000; Mystkowski et al., 2000; Morton et al., 2004).

The broad functional contributions of MCH cells may be attributed to their neurochemical diversity. MCH cells may utilize both classical neurotransmitters glutamate (Chee et al., 2015) or GABA (Jego et al., 2013) as well as other neuropeptides (Harthoorn et al., 2005; Mickelsen et al., 2017, 2019), and Mickelsen and colleagues (2019) recently identified two clusters of *Pmch* cells that could be distinguished based on their coexpression of *Cartpt* and *Tac3r*, the genes for cocaine- and amphetamine-regulated transcript (CART) and neurokinin 3 receptor (NK3R), respectively. However, functional differences between these clusters are not yet well defined.

CART is produced broadly throughout the brain and is also associated with varied behaviours (Koylu et al., 1997; Vrang, 2006; Subhedar et al., 2014). Within the hypothalamus, CART has been implicated in the control of multiple functions also associated with MCH, though they may have opposing effects. Notably, both peptides regulate energy balance, but while MCH mediates orexigenic actions that stimulate feeding in rats (Qu et al., 1996; Rossi et al., 1997; Clegg et al., 2003) and mice (Gomori et al., 2003; Glick et al., 2009), CART has shown both anorexigenic and orexigenic effects in a site-specific manner (Lau and Herzog, 2014; Subhedar et al., 2014). Intracerebroventricular administration of CART inhibited food intake (Kristensen et al., 1998; Aja et al., 2001) and reduced body weight (Larsen et al., 2000; Rohner-Jeanrenaud et al., 2002; Nakhate et al., 2010), but CART overexpression in specific hypothalamic areas, including the lateral, paraventricular, and ventromedial hypothalamic nucleus, increased food intake (Abbott et al., 2001; Kong et al., 2003; Smith et al., 2008; Farzi et al., 2018; Lau et al., 2018). Notably, half of MCH cells coexpress cocaine- and amphetamine-regulated transcript (CART; Broberger, 1999; Vrang et al., 1999; Elias et al., 2001; Cvetkovic et al., 2004; Croizier et al., 2010; Mickelsen et al., 2017, 2019; Wang et al., 2021). MCH and CART also both regulate sleep-wake behaviour and reproduction but in opposing directions. For example, while MCH cells are sleep-active (Hassani et al., 2009), and MCH promotes REM sleep (Verret et al., 2003), CART promotes wakefulness (Keating et al., 2010). To control reproductive behaviour, both MCH (Williamson-Hughes et al., 2005) and CART fibers (Leslie et al., 2001; Bogus-Nowakowska et al., 2011) contact neurons that produce gonadotrophin-releasing hormone, but MCH inhibits (Williamson-Hughes et al., 2005; Wu et al., 2009), while CART stimulates (Lebrethon et al., 2000; Parent et al., 2000; True et al., 2013), gonadotrophin-releasing hormone cells. Although the significance of MCH and CART coexpression is not known, their coexpression suggests a dual mechanism for acute, coordinated responses to internal or external changes in the environment (Kiss, 1988; Swanson, 1991). For this reason, it is possible that CART+ and CART− MCH cells may contribute to differential physiological roles.

MCH/CART+ cells are distinct from MCH/CART− cells in several ways. MCH/CART+ cells arise later in development than MCH/CART− cells (Brischoux et al., 2001; Cvetkovic et al., 2004; Risold et al., 2009; Croizier et al., 2010). They are anatomically divided, as MCH/CART+ cells are mostly distributed medially within the LHA, while MCH/CART− cells are predominantly in the lateral aspects of the LHA (Broberger, 1999; Vrang et al., 1999; Brischoux et al., 2001; Elias et al., 2001; Croizier et al., 2010; Wang et al., 2021). Furthermore, they have unique projection patterns (Brischoux et al., 2001; Cvetkovic et al., 2004; Risold et al., 2009), as MCH/CART+ cells predominantly project anteriorly towards the cerebral cortex, while MCH/CART− cells project posteriorly towards the spinal cord (Brischoux et al., 2001; Cvetkovic et al., 2004; Risold et al., 2009; Croizier et al., 2010). Finally, MCH/CART+ and MCH/CART− cells differentially express other neuropeptides. Notably, over two-thirds of MCH/CART+ cells coexpress NK3R, but this receptor is less abundant in MCH/CART− cells (Mickelsen et al., 2019; Fujita et al., 2021). Neurokinin B, the predominant ligand at NK3R (Steinhoff et al., 2014), is implicated in reproductive behaviours and physiology (Rance et al., 2010, 2013), and MCH/CART/NK3R+ cells could represent a further subdivision of the MCH/CART+ population.

As the MCH system is sexually dimorphic and the functional contributions of MCH/CART+ and MCH/CART− cells may be defined by their cellular properties, we described these two cell types in male and female mice based on their neuroanatomical distribution, electrical properties, and morphological characteristics. We generated high-resolution Nissl-based maps to chart the spatial distribution of MCH cells and their colocalization with CART and NK3R within the hypothalamus. Consistent with prior reports (Broberger, 1999; Vrang et al., 1999; Brischoux et al., 2001; Elias et al., 2001; Croizier et al., 2010; Wang et al., 2021), we also found that MCH/CART+ and MCH/CART− cells were most prevalent medial and lateral to the fornix, respectively. These two subpopulations also revealed different dendritic branching patterns, excitatory afferent inputs, and responses to current stimulation. Interestingly, while there were no overt sex differences in the distribution of MCH cell types, CART coexpression affected dendritic branching, cell resistance, and action potential firing depending on sex. Our findings indicated that in addition to differences in the neurochemical and spatial distribution patterns of CART+ and CART− MCH cells, they also presented distinctive electrical and morphological characteristics. These distinctions may thus underlie the unique contributions of each subpopulations to the overall scope of the MCH system.

### MATERIALS & METHODS

## Animals

All animal procedures were completed in accordance with guidelines and approval of the Animal Care Committee at Carleton University. Mice were group-housed at 21–22 ℃ with a 12:12 h light-dark cycle and provided with *ad libitum* access to water and standard mouse chow (Teklad Global Diets 2014, Envigo, Mississauga, Canada).

To visualize MCH neurons, *Pmch-cre* mice (Kong et al., 2010) were crossed to *R26-lox-STOP-lox-L10-Egfp* reporter mice (Krashes et al., 2014), kindly provided by Dr. B. Lowell (Beth Israel Deaconess Medical Center, Boston, MA), to generate *Mch-cre;L10-Egfp* mice expressing enhanced green fluorescent protein (EGFP) under the *Mch* promoter.

## Antibody characterization

**Table 1** lists the details of the following primary antibodies and how they were used in immunohistochemistry (IHC) or dual IHC and *in situ* hybridization experiments.

**Table 1.** List and details of primary antibodies.

| Antibody | Immunogen | Clonality, Isotype | Source, Product no., Lot no. | RRID | Titer |
| --- | --- | --- | --- | --- | --- |
| Rabbit anti-MCH | Full length MCH peptide | Polyclonal, IgG | Dr. E. Maratos-Flier | AB_2314774 | 1:2,000 |
| Rabbit anti-NK3R | Amino acids 375–452 at C-terminus of rat NK3R | Polyclonal, IgG | Novus Biologicals, NB300-102, F-4 | AB_10003270 | 1:150,000 |
| Rabbit anti-CART | Amino acids 61–102 of CART | Polyclonal, IgG | Phoenix Pharmaceuticals, H-003-61, 1625-3 | AB_2922768 | 1:2,000 |

**Rabbit anti-MCH antibody** was made and generously provided by Dr. E. Maratos-Flier (Beth Israel Deaconess Medical Center, Boston, MA). Antibody specificity was demonstrated by a lack of MCH-immunoreactivity in brain tissue from MCH knockout mice (Chee et al., 2013) and following MCH peptide absorption (Elias et al., 1998).

**Rabbit anti-NK3R antibody** was raised against amino acids 375–452 in the C-terminus of the rat NK3R protein. This antibody recognized a 50–52 kDa band from an immunoblot of rat brain homogenate (Le Brun et al., 2008). Our NK3R immunoreactivity pattern matched that reported in the rat hypothalamus (Ding et al., 1996; Burke et al., 2006), and there was a lack of staining following preabsorption with the NK3R peptide (Novus Biological, product datasheet).

**Rabbit anti-CART antibody** was raised against amino acids 61–102 of the CART protein. Antibody specificity was established by the lack of CART immunoreactivity when 1 ml of diluted antibody was preabsorbed with 10–100 μg of CART antigen (Arciszewski et al., 2009).

Secondary antibodies used were raised in donkey against the rabbit or were streptavidin-conjugated (**Table 2**).

**Table 2.** List and details of secondary antibodies.

| Antibody | Isotype | Source, Product no., Lot no. | RRID | Titer |
| --- | --- | --- | --- | --- |
| Donkey anti-rabbit Alexa Fluor 647 | IgG (H+L) | Thermo Fisher Scientific, A-31573, 2420695 | AB_2536183 | 1:500 |
| Biotin-SP AffiniPure donkey anti-rabbit | IgG (H+L) | Jackson ImmunoResearch, 711-065-152, 134054 | AB_2340593 | 1:500 |
| Cy3 Streptavidin |  | Jackson ImmunoResearch, 016-160-084, 146014 | AB_2337244 | 1:1,000<br>1:5,000 |

## Tissue preparation

*Mch-cre;L10-Egfp* mice (6–13 weeks) were anesthetized with an intraperitoneal (ip) injection of 7% chloral hydrate (700 mg/kg; Sigma-Aldrich, St-Louis, MO), and brain tissue was collected as previously described (Negishi et al., 2020). In brief, the mice were transcardially perfused with cold 0.9% saline and 10% formalin, and their brains were post-fixed in formalin for 12–24 h (4 °C) and cryoprotected in 20% sucrose for 12–24 h. Each brain was sliced along the coronal plane into five series of 30 μm-thick sections using a freezing stage microtome (Spencer Lens Co., Buffalo, NY) and collected in phosphate-buffered saline (PBS).

Free-floating tissue sections were stored in antifreeze solution at −20 °C for immunohistochemistry (IHC) experiments or were immediately mounted onto Superfrost Plus microscope slides (Fisher Scientific, Hampton, NH), air dried at room temperature (RT; 21–22 °C, 20 min) and −20 °C (30 min), then stored at −80 °C, as previously described (Bono et al., 2022) for Nissl staining. For fluorescence *in situ* hybridization (fISH) experiments, free-floating tissue sections were used immediately.

## Immunohistochemistry (IHC) and fluorescence in situ hybridization (fISH)

### Single-label IHC

As previously described (Chee et al., 2013; Negishi et al., 2020), free-floating slices were washed six times PBS for 5 min each (RT), blocked in 3% normal donkey serum (NDS; Jackson ImmunoResearch Laboratories, West Grove, PA; RRID: AB_2337258) prepared in PBS containing 0.05% sodium azide and 0.25% TritonX-100 (PBT-azide) for 2 h, and then incubated in rabbit anti-MCH antibody (1:2,000) overnight. After six 5-minute PBS washes (RT), the sections were incubated (2 h, RT) in donkey anti-rabbit Alexa Fluor 647 (1:500) and then rinsed with three 10-minute PBS washes (RT). The brain sections were mounted onto Superfrost Plus glass microscope slides (Fisher Scientific) and coverslipped with ProLong Diamond Antifade Mountant (Thermo Fisher Scientific) and 1.5 thickness glass (22-266-882P, Fisher Scientific).

### Dual-label IHC

Primary antibodies for anti-NK3R and anti-CART were both made in the rabbit (**Table 1**), thus, to minimize cross-reactivity between NK3R- and CART-immunoreactive (-ir) signals from rabbit-specific secondary antibodies, we labeled each antigen in series where labeling for NK3R immunoreactivity preceded that for CART.

Brain sections were rinsed via six exchanges of PBS (5 min each, RT), treated with 0.3% hydrogen peroxide (20 min), and rinsed with three PBS washes (10 min each, RT). The sections were blocked in 3% NDS (2 h) and immediately incubated (24 h, RT) with a rabbit anti-NK3R primary antibody (1:150,000), which was prepared in blocking solution. This anti-NK3R titer was established by serial dilution (Hoffman et al., 2016) and revealed that NK3R-ir signals were not detectable without tyramine signal amplification (TSA) (Hunyady et al., 1996; von Wasielewski et al., 1997; Tóth and Mezey, 2007).

After primary incubation, the sections were rinsed with six 5-minute PBS washes, then incubated in a biotinylated donkey anti-rabbit secondary antibody (1:500) for 1 h (RT) before rinsing again with three 10-minute PBS exchanges. The sections were then treated with an avidin-biotin-horseradish peroxidase solution (1:1:833; PK-6100, Vector Laboratories, Burlingame, CA; RRID: AB_2336819) for 30 min (RT), rinsed thrice with PBS (10 min each), and incubated (20 min) in a PBT solution comprising 0.005% hydrogen peroxide and 0.5% biotinylated (EZ-Link sulfo-NHS-LC-biotin; ThermoFisher Scientific) tyramine (Sigma-Aldrich, St. Louis, MO), which was made in-house (Diniz et al., 2020). The sections were rinsed with three 10-min PBS exchanges and treated with Cyanine 3 (Cy3)-conjugated streptavidin (1:1,000) in 3% NDS without azide for 2 h (RT).

To prepare for anti-CART immunolabeling, the sections were washed in three 10-minute PBS washes and incubated in the rabbit anti-CART antibody (1:2,000) for 24 h (RT). They were then rinsed six times in PBS (5 min each) before treatment with donkey anti-rabbit Alexa Fluor 647 antibody (1:500) for 2 h (RT). Following three final PBS washes (10 min each), the slices were promptly mounted and coverslipped with ProLong Diamond Antifade Mountant (ThermoFisher Scientific).

### Dual IHC and fISH

Immediately after tissue sectioning, free-floating tissues were rinsed with six 5-minute PBS washes (RT), blocked in 3% NDS for 1 h, and then incubated with a rabbit anti-MCH primary antibody (1:2,000) for 2 h. Subsequently, the sections were washed in three 5-minute PBS exchanges (RT), treated with donkey anti-rabbit Alexa Fluor 647 secondary antibody (1:500) for 2 h, washed again in three 5-minute PBS exchanges, and then mounted onto Superfrost Plus microscope slides (Fisher Scientific). The slides were air dried at RT (X min) and –20 °C (30 min), and then sealed in a microscope slides box (HS15994GF, Fisher Scientific) for storage at −80 °C to promote tissue adherence.

One week later, the slides were warmed in a HybEZ II oven (Advanced Cell Diagnostics (ACD), Newark, CA) at 37 °C for 45 min, dehydrated in ascending ethanol concentrations (50%, 70%, and 100%) at RT for 5 min each, and air-dried at RT for 15 min before implementing RNAscope-mediated fISH as previously described (Bono et al., 2022). Three parallel sets of slides were established for RNAscope fISH to determine the coexpression of *Pmch* and *Egfp* mRNA; hybridization for mouse peptidylprolyl isomerase B (*Ppib*) to verify RNA and tissue quality; and hybridization for *Bacillus* dihydrodipicolinate reductase (*dapB*) to assess background staining. All RNAscope probes were acquired from ACD and detailed in **Table 3**.

**Table 3.**
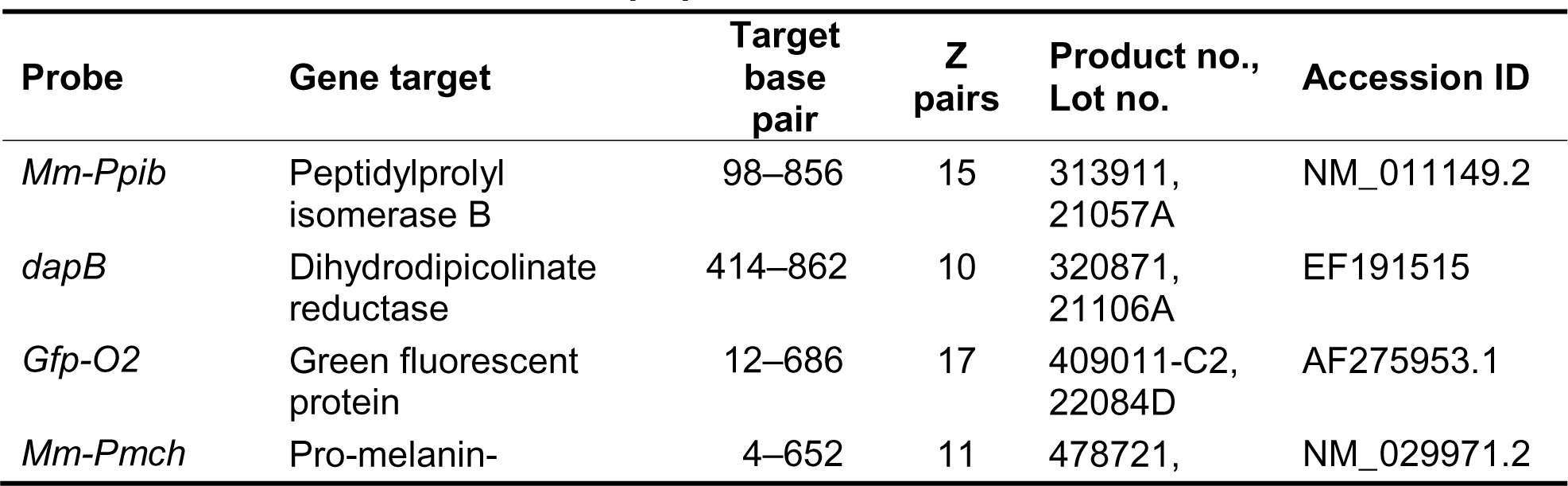
List and details of RNAscope probes.

Briefly, the sections were treated with hydrogen peroxide (322335, ACD) for 10 min (RT) to quench endogenous peroxidase activity and rinsed twice for 3 min each in distilled water (RT). The slides were then briefly (30 s) acclimated in distilled water at 99 °C before incubated with Target Retrieval Reagent (322000, ACD) for 5 min (99 °C). The slides were again rinsed in distilled water (15 s, RT), submerged in 100% ethanol (3 min, RT), and allowed to air-dry for 15 min (RT). A hydrophobic barrier was drawn around the tissue sections to prevent subsequent reagents from running off the slide.

The tissue was treated with Protease III (322337, ACD) at 40 °C for 30 min and washed with distilled water (3 min each, RT). RNAscope probes for *Pmch* and *Egfp*, *Ppib*, and *dapB* were applied to their respective slides and to hybridize for 2 h at 40 °C before washing three times (5 min each, RT) with Wash Buffer (3100931, ACD). Probe signals were amplified by the following incubations at 40 °C: AMP 1 (30 min; 323101, ACD), AMP 2 (30 min; 323102, ACD), and AMP 3 (15 min; 323103, ACD) that alternated with two exchanges in fresh Wash Buffer (5 min each, RT).

All tissues were treated with HRP-C1 (15 min, 40 °C; 323104, ACD), washed twice in Wash Buffer (5 min each, RT), and incubated with TSA Plus Cyanine 3 (1:750; 30 min, 40 °C; NEL744E001KT, PerkinElmer, Waltham, MA) prepared in TSA Buffer (322809, ACD). After two washes in Wash Buffer (5 min each, RT), the tissues were incubated in HRP Blocker (15 min, 40 °C; 323107, ACD). HRP-C2 (15 min, 40 °C; 323105, ACD) was then applied to the tissue followed by Opal 520 (1:500, 30 min, 40 °C; FP1487001KT, Akoya Biosciences, Marlborough, MA) prepared in TSA Buffer. The tissue was rinsed twice with Wash Buffer (5 min each, RT), incubated with HRP Blocker (15 min, 40 °C), and again rinsed in Wash Buffer (5 min each, RT). All slides were coverslipped with ProLong Diamond Antifade Mountant (ThermoFisher Scientific).

## Nissl stain

Nissl stains (Simmons and Swanson, 1993) were performed on tissues adjacent to our experimental tissue to determine the levels and cytoarchitectural boundaries within each slice. As previously described (Negishi et al., 2020), the mounted tissues were dehydrated in serial washes of 50%, 70%, 95%, and three exchanges of 100% ethanol (3 min each, RT) and then delipidated in two incubations with fresh xylene for 10 min and 15 min, respectively. The tissues were then rehydrated by six ethanol washes (3 min each) repeated in reverse order and then soaked in distilled water for 3 min.

The tissues were stained with a 0.25% w/v thionine dye solution (pH 4.5) at RT for about 5 s. If the staining appeared too dark, the slides were rinsed in 4% glacial acetic acid to remove excess dye (or in distilled water if dye is excessive), and if the staining was too faint, the slides were dipped again in thionine solution until the grey and white matter were clearly distinguishable by eye. The reaction was quenched in distilled water, and the slides were dehydrated in increasing concentrations of ethanol (3 min each), cleared in in two xylene incubations (15 min), and then coverslipped using Richard-Allan Scientific Mounting Media (Fisher Scientific).

## Microscopy

All images were captured using an Eclipse Ti2 inverted microscope (Nikon Instruments Inc., Mississauga, Canada) equipped with a motorized stage, DS-Ri2 color camera (Nikon), Prime 95B CMOS camera (Photometrics, Tucson, AZ), and Plan Achromat 10× (0.45 numerical aperture), 20× (0.75 numerical aperture), or 40× (0.95 numerical aperture) objective lenses. Captured images were stitched and/or adjusted for brightness using NIS Elements software (Nikon), exported as TIFF files, and then imported into Adobe Illustrator 26.2.1 (Adobe Systems Inc., San Jose, CA) for analysis and figure assembly.

### Epifluorescence imaging

Large overview images of the entire brain slice in fluorescence were acquired with a Prime 95B CMOS camera using the 10× objective lens. Tissue structures were illuminated by a 561-nm wavelength light via a SPECTRA X engine (Lumencor, Beaverton, OR). Tiled images were stitched into a single overview image of the entire slice using NIS Elements software.

### Confocal imaging

High resolution images (2,048 × 2,048 pixels) of the hypothalamus were captured with a C2 confocal system (Nikon) using either a 10× or 20× objective lens. Regions of interest were selected from overview images (section 2.7.1) and acquired with a 488-, 561-, or 640-nm wavelength lasers to visualize native EGFP or Opal 520, Cy3 or Alexa Fluor 594, and Alexa Fluor 647, respectively. Resulting images were stitched to simultaneously display individual cells and an overview of the entire hypothalamus, where applicable, and pseudo-colored green, orange, or magenta, respectively.

In RNAscope experiments, the laser and image settings were adjusted to eliminate visible fluorescence in the *dapB* negative control sections and then applied to capture and process images from experimental tissue. All images were exported as TIFF files to Adobe Illustrator for assembly and quantification.

### Brightfield imaging

Large field-of-view images of the entire brain slice from Nissl-stained tissues were acquired with the DS-Ri2 color camera using a 10× objective lens and were stitched using NIS Elements (Nikon) to resolve white and gray matter distinctions within each slice. These were exported as TIFF files and imported into Adobe Illustrator for plane-of-section analysis and alignment with epifluorescence or confocal images.

## Image analysis

### Nissl-based parcellation

Stitched overview brightfield and epifluorescence images of each entire brain slice were imported into Adobe Illustrator and were overlayed and aligned to one another to determine its corresponding *Allen Reference Atlas* (*ARA*) level (Dong, 2008). Nissl-based parcellations were drawn using the *Pen* tool to distinguish the boundaries of each brain region or white matter. Parcellations were defined based on cell morphology, density, and directionality, and they were superimposed onto the matching epifluorescence overview image of the brain slice. Additionally, higher magnification confocal images were aligned to the epifluorescence overview image thus effectively transferring parcellations onto confocal images also.

### **Plane-of-section analysis and *ARA* level assignment**

If the hypothalamus within a brain slice was uneven dorsoventrally, it was subdivided into three horizontal segments, which were demarcated as the horizontal region between the mamillary bodies dorsally to the ventral tip of the globus pallidus (dorsal segment); then to the fornix ventrally (middle segment); and then to the ventral edge of the hypothalamus slice (ventral segment). A slice was only included for cell quantification if: i) the same level was assigned to the top two horizontal segments, which comprised the majority of MCH cells, and ii) if a matching level was found for the bottommost horizontal segment on another slice within the same brain series.

### Mediolateral division of the hypothalamus

To quantify the cellular distribution mediolaterally, each *ARA* map was divided into four 0.5 mm-wide vertical sections encompassing the 3^rd^ ventricle to the lateral border of the hypothalamus. We quantified the proportion of every cell type expressed within each vertical segment to determine their mediolateral distribution.

## Cell quantification and atlas-based mapping

An epifluorescence overview image of the entire brain slice and a high-magnification confocal image comprising preselected regions of interest were assembled in an Adobe Illustrator file to be aligned so that both images together would simultaneously display the entire brain slice and the distribution of cells within our region of interest. Each cell type was marked with a filled circle using the *Ellipse* tool and then counted by determining the number of circles selected in the *Document Info* menu of Adobe Illustrator. The relative positions of filled circles were mapped onto *ARA* brain templates (Dong, 2008). Mean cell counts per level was reported unilaterally. Total cell counts per brain was reported bilaterally.

Number of cells reported was corrected for oversampling, as previously described (Chee et al., 2013; Negishi et al., 2020; Bono et al., 2022), using the Abercrombie formula (Abercrombie, 1946): 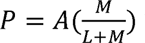 where *P* is the corrected count, *A* is the original count, *M* is the mean tissue thickness (17.42 μm) determined from five brain slices, and *L* is the mean cell diameter (14.60 μm) determined from 107 MCH-ir cells.

## Electrophysiology

### Slice preparation

Male and female *Mch-cre;L10-Egfp* mice (4–23 weeks) were anesthetized with 7% chloral hydrate (700 mg/kg, ip) and then were transcardially perfused with ice-cold N- methyl-D-glucamine based (NMDG)-slice artificial cerebrospinal fluid (ACSF) containing (in mM): 50 NMDG, 1.25 KCl, 10 HEPES, 10 MgSO_4_, 12.5 glucose, 15 NaHCO_3_, 0.25 CaCl_2_, 1 thiourea, 2.5 L-ascorbic acid, 1.5 sodium pyruvate (300 mOsm/L, pH 7.4). The brains were rapidly removed from the skull and were sliced with a vibratome (VT1000S, Leica, Wetzlar, Germany) into 250 μm-thick coronal sections. Slices were incubated in NMDG-slice ACSF at 37 °C for 5 min and then transferred to bath ACSF comprising (in mM): 124 NaCl, 3 KCl, 1.3 MgSO4, 1.4 NaH_2_PO_4_, 10 glucose, 26 NaHCO_3_, 2.5 CaCl_2_ (300 mOsm/L) for 5 min (37 °C). Slices were allowed to recover at RT for at least one hour prior to slice recording. All slice and bath ACSF solutions used were constantly carbogenated (95% O_2_, 5% CO_2_).

### Whole-cell patch-clamp recordings

Slices were equilibrated in a recording chamber and maintained by a continuous flow (1–1.5 ml/min) of carbonated bath ACSF at 31–32 ℃. MCH cells were identified by epifluorescence illumination of native EGFP and visualized by infrared differential interference contrast at 40× magnification on either an Eclipse FN1 microscope (Nikon) equipped with a mercury lamp (C-SHG1, Nikon), pco.panda 4.2 sCMOS camera (Excelitas PCO GmbH, Kelheim, Germany), and NIS Elements imaging software (Nikon) or an Examiner.A1 microscope (Carl Zeiss Inc, Oberkochen, Germany) equipped with a mercury lamp (HXP 120, Zeiss), an AxioCam Mrm camera (Zeiss), and AxioVision 4.8 software (Zeiss).

Glass micropipettes used for recording were pulled using a Flaming/Brown Micropipette puller (P-1000, Sutter Instruments, Novato, CA) with a pipette resistance of 7–10 MΩ when backfilled with an internal solution containing (in mM): 120 K-gluconate, 10 KCl, 10 HEPES, 1 MgCl_2_, 1 EGTA, 4 MgATP, 0.5 NaGTP, 10 phosphocreatine, and 0.2% biocytin (pH 7.24, 280– 290 mOsm/L). All recordings were acquired using a MultiClamp 700B amplifier (Molecular Devices, San Jose, CA) and digitized by either a Digidata 1440A or 1550B (Molecular Devices) running pClamp 10.3 or 11.2 software (Molecular Devices).

### Electrophysiological parameters

All recordings were filtered at 1 kHz, and all reported values were corrected for a +15 mV liquid junction potential. In-between recordings, the cells were held at a holding potential (V_h_) = −60 mV. Electrophysiological data was collected using Clampfit 10.3 or 11.2 software (Molecular Devices), unless indicated otherwise.

**Capacitance** was calculated from the average of five −10 pA hyperpolarizing current steps using a formula for non-isopotential cells (Golowasch et al., 2009). Capacitance values (C_m_) were calculated as 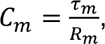, where *τ*_m_is the time constant determined by fitting the voltage curve obtained in response to each hyperpolarizing step and R_m_ is the membrane resistance. Curve fitting was performed with two exponentials, using the Levenberg-Marquardt algorithm in Clampfit 11, where the slowest *τ* term corresponded to the charging of the membrane capacitance.

**Input resistance** was determined by measuring the voltage output from a −30 pA hyperpolarizing current step (3 s).

### Current–voltage (I–V) relationship

Current change was determined in voltage-clamp at V_h_ = −60 mV in response to −10 mV voltage steps (250 ms) elicited from −40 mV to−110 mV.

**Resting membrane potential (RMP)** was averaged over two minutes in current-clamp mode with no current injection immediately after establishing whole-cell configuration.

**Cell excitability** was determined by rheobase, and the number of action potentials elicited from V_h_ = −60 mV by +10 pA depolarizing current steps (3 s) evoked from 0 pA to 90 pA. Action potentials were considered if their peak amplitudes exceeded 0 mV.

**Excitatory synaptic input** was measured as the spontaneous excitatory post-synaptic current (sEPSC) events recorded at V_h_ = −60 mV over a 5-min period. sEPSC frequency, interevent intervals, and amplitude were detected using MiniAnalysis (Synaptosoft, Decatur, GA). As sEPSC recordings were collected in the absence of GABA_A_ receptor antagonists or blockers, it is possible that sEPSC events may be contaminated by chloride-mediated events. However, as the equilibrium potential of chloride in our recording conditions (E_Cl_ = −62.8 mV) was negative to the holding potential, any inhibitory chloride-mediated events would emerge as positive currents, and we restricted sEPSC analyses to only negative current events. All or up to 70 events recorded from each cell were included in the analyzed dataset. The distributions of sEPSC amplitudes and areas were normalized and fitted by the amplitude version of the Gaussian peak function (GaussAmp) in Origin 2018b (OriginLab Corporation, Northampton, MA). To determine the number of sEPSC subpopulations detected, the cumulative distribution curve was fitted with increasing number of peaks until the goodness of fit (*R^2^*) was unchanged. Each fit underwent repeated iterations until the chi-square per degrees of freedom was unchanged (Bono et al., 2022). sEPSC kinetics were determined by the rise time, which was calculated as the slope at 10–90% of each sEPSC rising phase, and decay time, which was calculated as *τ* based on a single-exponential fit from the peak to the end of the sEPSC decay phase. Amount of charge carried by each sEPSC was determined by the area under the curve of each event. Cells were excluded if noise threshold ≥ 9, if the cell was identified as an outlier, in at least 2 parameters, by ROUT test when Q = 0.1%.

### ***Post hoc* labeling of *Mch-cre;L10-Egfp* cells**

All recordings were maintained in whole-cell configuration for at least 30 min to ensure full penetration of biocytin fill throughout the recorded cell. At the end of the recording, the pipette was gently extracted by slowly gliding the pipette in a diagonal along the XZ plane, as described by Swietek et al. (2016).

### Thick tissue IHC

The whole slice was immediately submerged in 10% formalin for 15–18 h and then rinsed in six 5-min PBS washes (RT). The slices were permeabilized by incubation in PBS comprising 2% TritonX-100 for 45 min and blocked in 3% NDS prepared in PBT-azide composed of 0.25% TritonX-100 and 0.02% sodium azide for 2 h. Next, slices were incubated with rabbit anti-CART primary antibody (1:2,000) for 48 h at 4 ℃.

The slices were rinsed in six 5-min exchanges of PBS (RT) and then incubated in donkey anti-rabbit Alexa Fluor 647 (1:1,000) and Cy3-conjugated streptavidin (1:5,000; Jackson ImmunoResearch; AB_2337244) for 2 h (RT) prepared in 3% NDS. Slices were wet-mounted onto Superfrost Plus microscope slides for confocal imaging with a 20× objective lens to determine native EGFP and CART-ir signals in biocytin-filled cells. Only biocytin-filled cells that expressed EGFP were included in our analyses.

### 3,3’-diaminobenzidine (DAB) labeling of biocytin-filled cells

Following imaging analysis, the slides were soaked in PBS at RT and gently slipped off the microscope slide. The free-floating brain slices were prepared for morphological analysis. They were treated with 1% hydrogen peroxide in PBT for 30 min (RT) to quench endogenous peroxidase activity, rinsed in three 10- min PBS exchanges (RT), and treated to form the avidin-biotin complex in PBS (1:1:500; Vector Laboratories; AB_2336819) for 72 h (4 ℃). After three PBS washes (10 min each), the slices were incubated in a nickel-enhanced DAB solution, which was prepared according to manufacturer instructions, until the cells were dark black, and the cell outline was clearly visible (∼5 min). The thick slices were then mounted onto Superfrost Plus microscope slides, dehydrated in increasing ethanol concentrations (50%, 70%, 95%, and 100%) for 6 min each, delipidated in 100% xylene for 2 h, and coverslipped with Richard-Allan Scientific Mounting Media. Morphological features of the biocytin-filled DAB-stained cell was matched via brightfield imaging using a 20× objective lens.

## Morphological reconstruction and analysis

For cell reconstruction, the thick tissue slice was visualized in brightfield on a BX51 (Olympus, Tokyo, Japan) upright microscope that was equipped with a CX9000 camera (MBF Bioscience, Williston, VT) and fully motorized stage. Cells were analyzed live using an oil immersion Olympus Plan N 100× objective lens (0.95 numerical aperture). The soma, dendrites, and axon of each cell were traced using Neurolucida 2020.1.3 software (MBF Bioscience). A single line with varying widths was traced through the center of each process to match the thickness of each axon and/or dendrite.

Each reconstructed cell was exported into Neurolucida Explorer 2019.2.1 (MBF Bioscience). The number of nodes was determined by branched structure analysis, and the total length of each process and distance of each node from the soma were determined by Sholl analysis (Sholl, 1953), with each concentric ring increasing by a 10 μm radius. Primary dendrites were defined as those originating at the soma; secondary dendrites branched from primary dendrites, and tertiary dendrites branched from secondary dendrites. Similarly, primary nodes originated on primary dendrites, while secondary nodes originated on secondary dendrites, and henceforth. Cell tracings were exported as vectors and imported into Adobe Illustrator.

## Statistics

Data are represented as mean ± SEM, and the sample size, shown in parenthesis within the figures, is represented as *n* for number of cells and *N* for number of mice. Means compared across time, voltage, distance, or node order were compared using two-way, where †, p < 0.05; ††, p < 0.01; †††, p < 0.001 or mixed-design (if some data points were missing) ANOVA, where ^, p < 0.05 with the Geisser-Greenhouse correction for lack of sphericity. If there were significant main effects or interaction, a Bonferoni multiple comparison post hoc test was also implemented, where significance was determined as: *, *p* < 0.05; **, *p* < 0.01; ***, *p* < 0.001. Three-way ANOVA was used to test for sex differences between cell types across time, current injection, voltage range, or node order. Tests were chosen based on normality and homogeneity of variance. All statistical analyses were performed in Prism 9.3.1 (GraphPad Software Inc., San Diego, CA). Graphs and sample electrophysiological recordings traces were obtained using Prism and OriginPro (OriginLab, Northampton, MA), respectively.

### RESULTS

## Validity and specificity of EGFP*^Mch^* expression in *Mch-cre* mice

We selected MCH neurons by native EGFP fluorescence (EGFP*^Mch^*) in *Mch-cre;L10-Egfp* mice. EGFP*^Mch^* was detected only within the hypothalamus (**Figure 1a, b**), and nearly all MCH-ir neurons (92 ± 2%, N = 5) expressed EGFP*^Mch^* (**Figure 1c**). Importantly, we also determined the specificity of EGFP*^Mch^* labeling for MCH neurons and found that 81 ± 4% (N = 5) of EGFP*^Mch^* cells coexpressed MCH immunoreactivity overall (**Figure 1d**). However, by analyzing the distribution of MCH-ir and EGFP*^Mch^* signals across the MCH field rostrocaudally, we found that MCH-negative EGFP*^Mch^*cells were largely restricted to *ARA* level (L) 73 (**Figure 1e**; N = 4).

**Figure 1.**
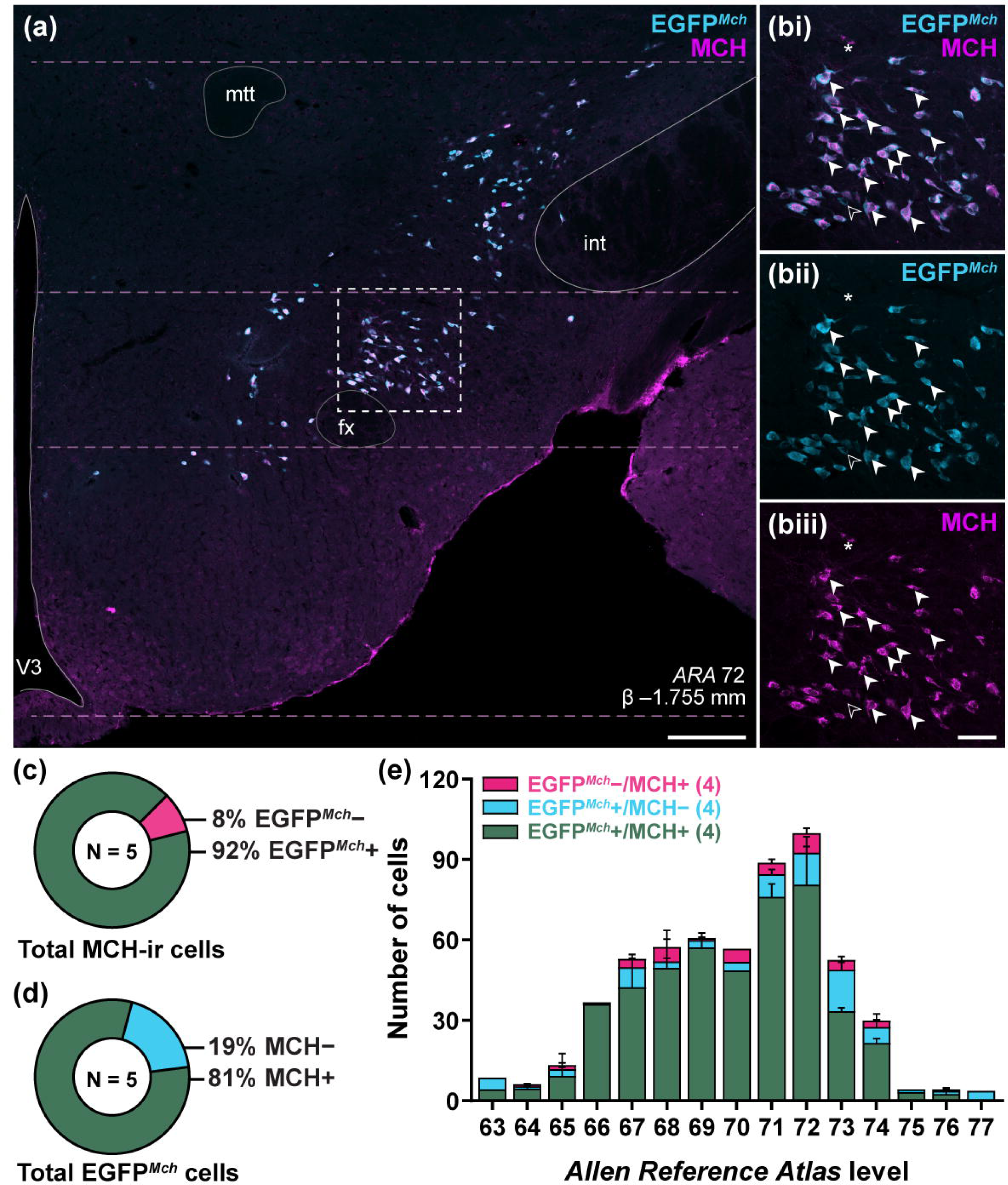
Validation of hypothalamic EGFP*^Mch^* expression in *Mch-cre* mice. Representative merged channel confocal photomicrographs from level 72 of the *Allen Reference Atlas* (*ARA*; Dong, 2008) at −1.755 mm relative to Bregma β within the hypothalamus (***a***). Merged channel confocal photomicrograph from outlined area in ***a*** (***bi***) of native EGFP fluorescence (EGFP*^Mch^*; ***bii***) and MCH immunoreactivity (***biii***) from the brain of a *Mch-cre;L10-Egfp* mouse. Representative sample of EGFP*^Mch^*cells that coexpressed MCH immunoreactivity (white arrowheads), EGFP*^Mch^*cells that were MCH-negative (open arrowheads), and MCH- immunoreactive (MCH-ir) cells that did not express EGFP*^Mch^* (* asterisk; ***b***). Percentage of MCH-ir cells marked by the presence (dark teal) or absence (magenta) of EGFP*^Mch^* (***c***). Percentage of EGFP*^Mch^*-labeled cells marked by the presence (dark teal) or absence (light blue) of MCH-ir signals (***d***). Anteroposterior distribution of the mean number of EGFP*^Mch^*−/MCH+ cells (magenta), EGFP*^Mch^*+/MCH− cells (light blue), and dual-labeled EGFP*^Mch^*+/MCH+ cells (dark teal) at each level of the hypothalamus assigned in accordance with the *ARA*. Only brain slices in which the dorsal and middle segment (demarcated by dashed pink lines, in ***a***) corresponded to the same ARA level were included in our dataset. Scale bar: 200 μm (***a***); 50 μm (***b***). fx, fornix; int, internal capsule; mtt, mammillothalamic tract; V3, third ventricle.

In the anterior MCH field (L63–66), we observed scattered MCH-ir cells that did not express EGFP (EGFP*^Mch^*−/MCH+) and EGFP-labeled cells that did not express MCH immunoreactivity (EGFP*^Mch^*+/MCH−) in the medial zona incerta (ZI) and the posterior part of the anterior hypothalamic nucleus (AHNp; **Figure 2a–c**). Towards the middle of the MCH field (L67–72), we saw robust colocalization between EGFP*^Mch^* and MCH immunoreactivity (EGFP*^Mch^*+/MCH+), with only few non-colocalized cells in the ZI, LHA, and anterior part of the dorsomedial hypothalamic nucleus (DMHa), parasubthalamic nucleus (PSTN) and subthalamic nucleus (STN; **Figure 2d–i**). However, at L73, we consistently observed an EGFP*^Mch^*+/MCH− cluster that appeared immediately dorsal to the fornix (**Figure 2j**), as described by Beekly et al. (2020). In the posterior hypothalamic MCH field (L74–77), we found consistent EGFP*^Mch^+*/MCH+ colocalization, with only a few EGFP*^Mch^*−/MCH+ cells and a few EGFP*^Mch^*+/MCH− cells in the medial LHA (**Figure 2k, l**).

**Figure 2.**
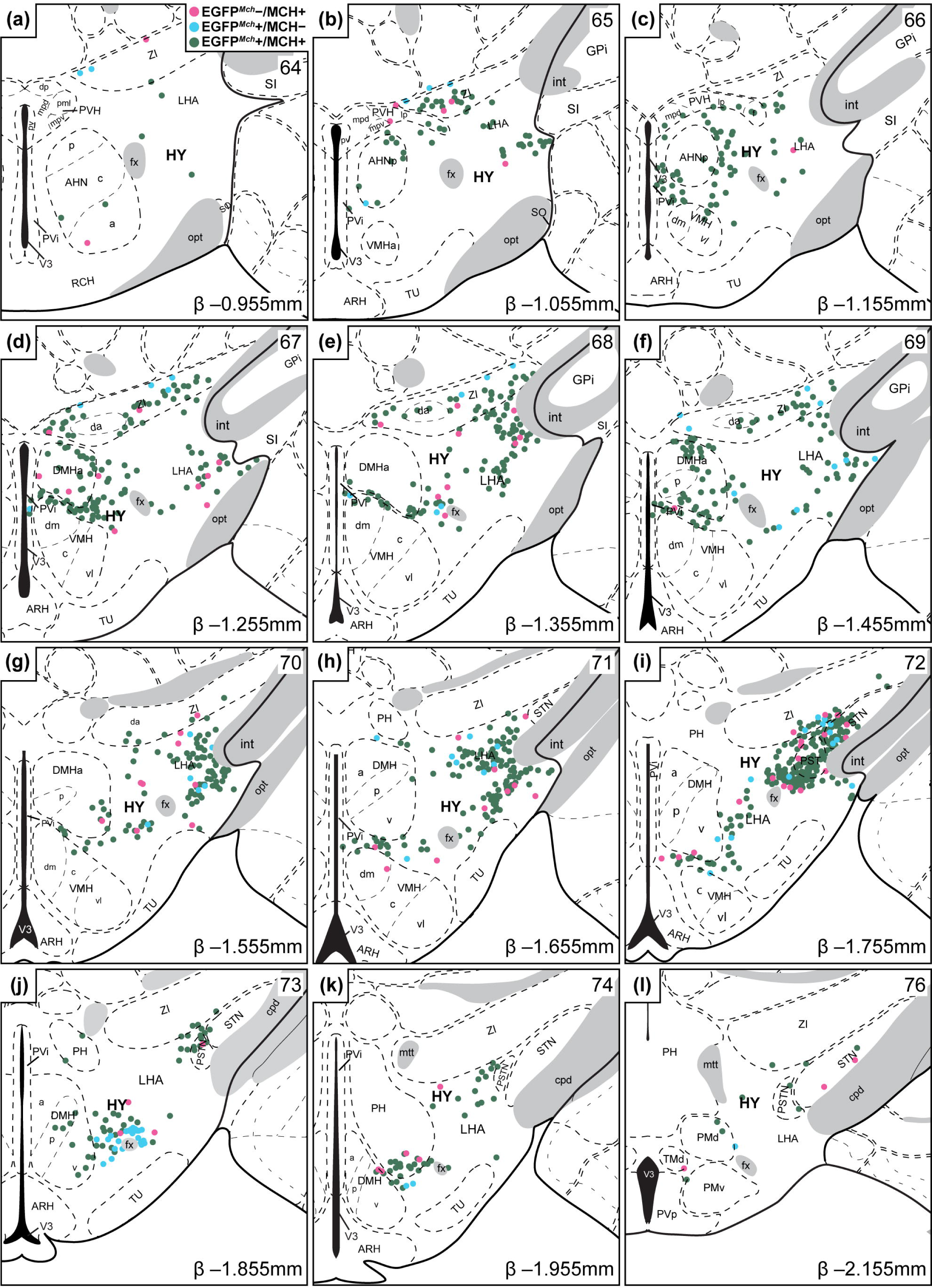
Spatial distribution of EGFP*^Mch^* expression in *Mch-cre* mice. Representative coronal maps using *Allen Reference Atlas* (*ARA*; Dong, 2008) brain templates arranged from rostral to caudal (***a*–*l***) showing the relative distribution of EGFP*^Mch^*−/MCH+ cells (magenta circles), EGFP*^Mch^*+/MCH− cells (light blue circles), and dual-labeled EGFP*^Mch^*+/MCH+ cells (dark teal circles) from the hypothalamus of *Mch-cre;L10-Egfp* mice. Each panel includes the corresponding *ARA* level (top right), stereotaxic coordinate inferred from Bregma (β; bottom right), and brain region labels according to *ARA* nomenclature.

Overall, we found that EGFP*^Mch^* cells robustly identified MCH neurons in *Mch-cre;L10-Egfp* mice throughout the MCH field, with the exception of a distinctive cluster of EGFP*^Mch^*+/MCH− cells situated within or around the fornix though they may appear smaller and fainter than surrounding MCH-ir cells (**Figure 3a, b**). We performed dual *in situ* hybridization for *Gfp* mRNA and *Pmch* mRNA, but nearly all EGFP*^Mch^*+/MCH− cells did not express either transcript (**Figure 3c**) and may thus reflect legacy EGFP expression in cells where the *Mch* promoter is no longer active. However, though rare, an EGFP*^Mch^* cell that was not MCH-ir can express *Pmch* (**Figure 3ciii**, asterisk).

**Figure 3.**
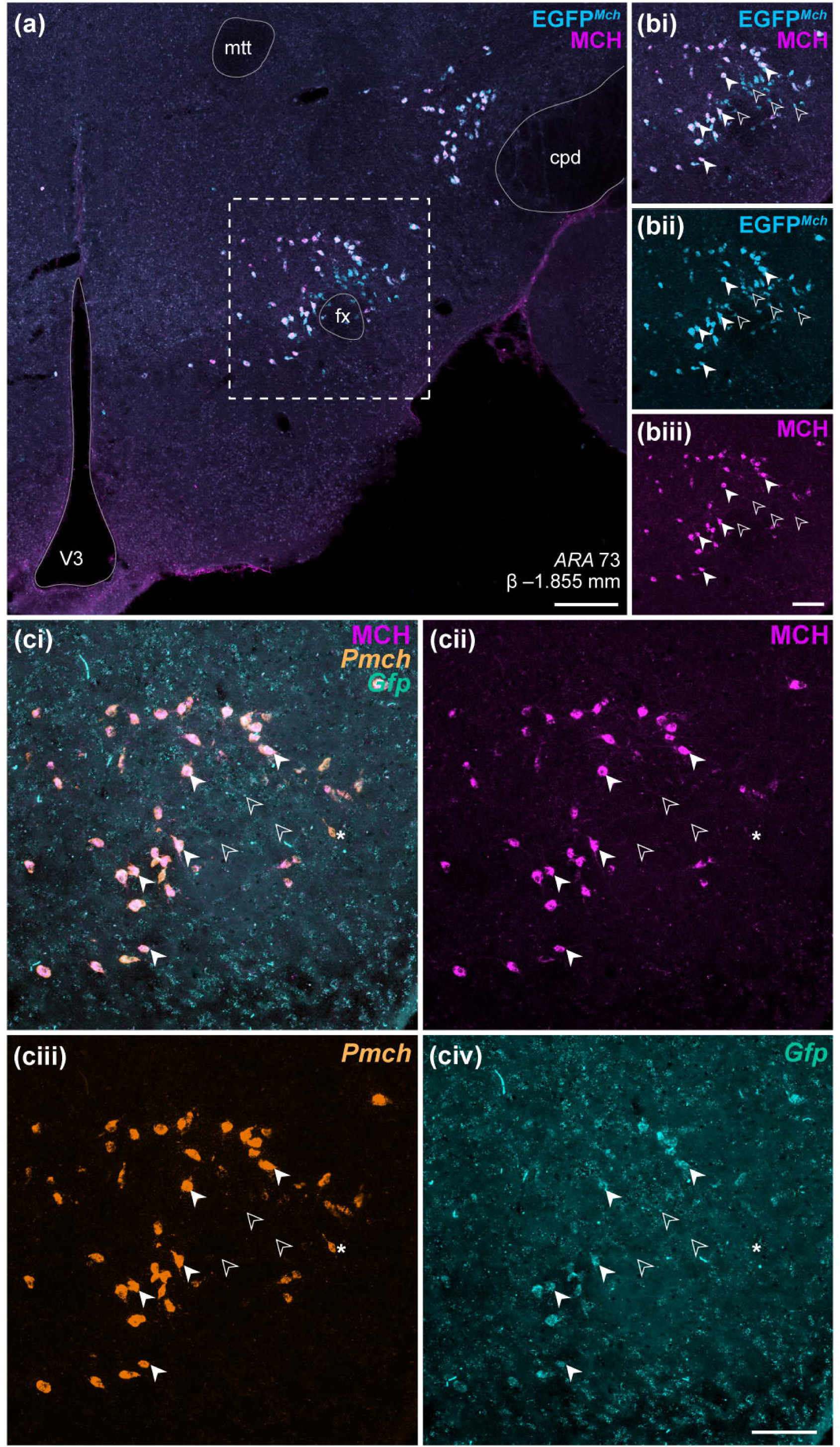
Absence of MCH immunoreactivity, *Pmch* mRNA, or *Gfp* mRNA at perifornical EGFP*^Mch^*cells. Merged channel low (***a***) and high magnification (***bi***) confocal photomicrographs from outlined area around the fornix (fx) in ***a*** of native EGFP fluorescence (EGFP*^Mch^*; ***bii***) and MCH immunoreactivity (***biii***) within the hypothalamus of *Mch-cre;L10-Egfp* mice. Representative sample of EGFP*^Mch^* cells that were MCH-immunoreactive (white arrowheads) or that were MCH- negative (open arrowheads). Merged channel confocal photomicrograph (***ci***) of MCH immunoreactivity (***cii***), *Pmch* mRNA hybridization (***ciii***), and *Gfp* mRNA hybridization (***civ***) captured from the same cells and area outlined in ***a*** following dual-label RNAscope *in situ* hybridization. Representative sample of EGFP*^Mch^* cells (***ci****–**iv***) previously identified in ***b*** that were MCH-ir and expressed both *Pmch* and *Gfp* hybridization (filled arrowheads), that did not express MCH-ir but expressed *Pmch* (* asterisk), or that did not express MCH-ir, *Pmch*, or *Gfp* hybridization (open arrowheads). Scale: 200 μm (***a***); 100 μm (***b*, *c***). cpd, cerebral peduncle; fx, fornix; mtt, mammillothalamic tract; V3, third ventricle.

## Spatial distribution of CART and/or NK3R coexpression at MCH neurons

EGFP*^Mch^* expression robustly represented MCH neurons (**Figure 1, 2**), thus we determined the colocalization of CART and NK3R immunoreactivity in EGFP*^Mch^* cells (**Figure 4a**). The number and proportion of each EGFP*^Mch^* cell type was similar between female and male mice. EGFP*^Mch^*- only cells comprised over half of all EGFP*^Mch^* cells counted from the hypothalamus of female (53 ± 7%, N = 4; **Figure 4b**) and male (55 ± 2%, N = 2; **Figure 4c**) *Mch-cre;L10-Egfp* mice. There were relatively equal proportions of EGFP*^Mch^* cells that coexpressed both CART and NK3R (EGFP*^Mch^*/CART/NK3R; female: 21 ± 3%; male: 23 ± 3%) or that coexpressed CART only (EGFP*^Mch^*/CART; female: 22 ± 4%; male: 20 ± 2%), thus nearly half of all EGFP*^Mch^* cells were CART-positive. The remaining EGFP*^Mch^* cells included a small proportion of EGFP*^Mch^* cells that coexpressed NK3R only (EGFP*^Mch^*/NK3R; female: 4 ± 1%; male: 2 ± 0%). All four subpopulations of EGFP*^Mch^* cells could be seen throughout the rostrocaudal axis of the hypothalamus (L63–L77), but the availability of EGFP*^Mch^*cells gradually peaked at L71 then decreased posteriorly in a normally distributed pattern (**Figure 4d**).

**Figure 4.**
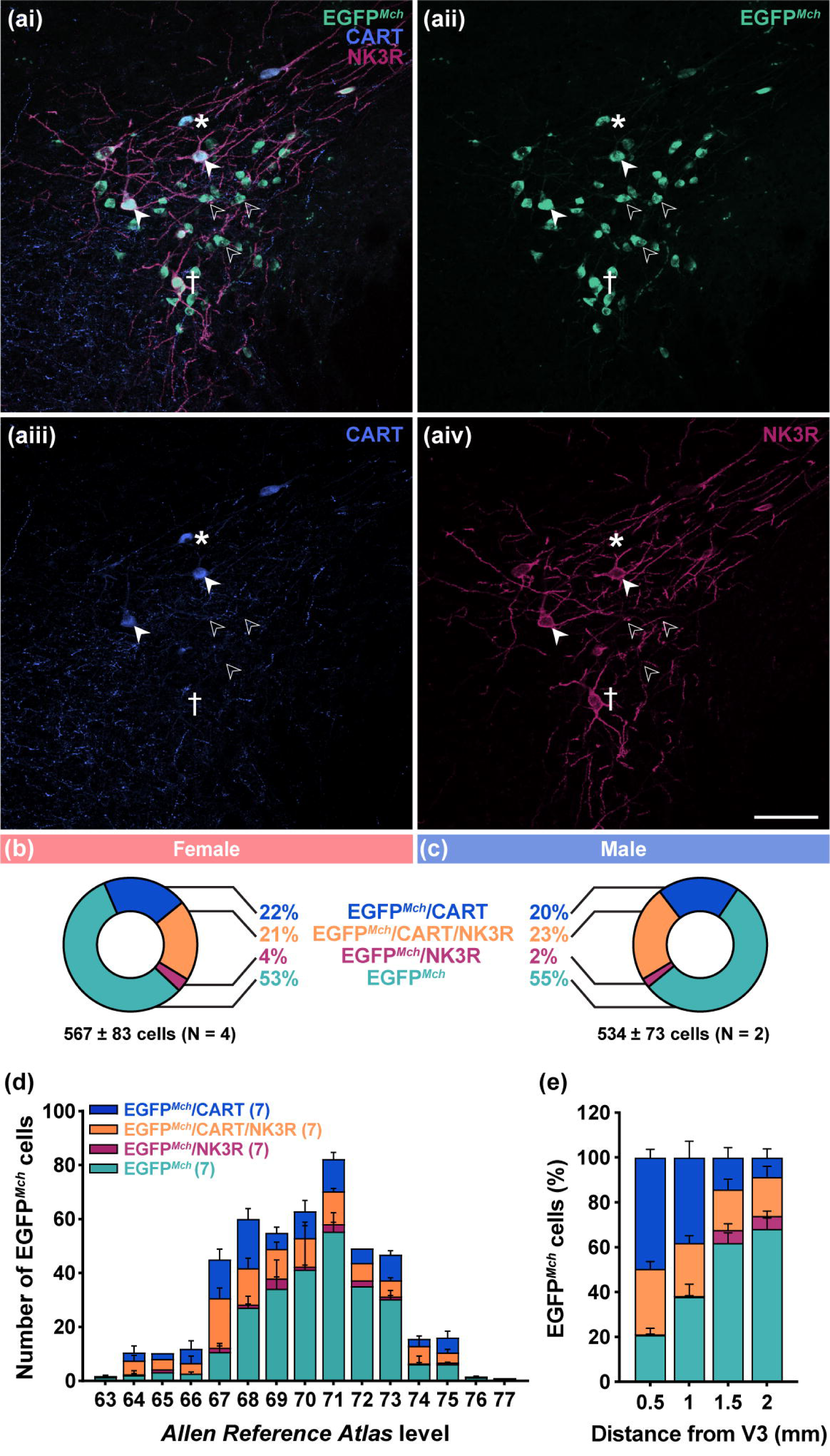
Diverse MCH cell types defined by CART and NK3R coexpression. Merged channel (***ai***) confocal photomicrographs of native EGFP fluorescence (EGFP*^Mch^*; ***aii***), CART immunoreactivity (***aiii***), and NK3R immunoreactivity (***aiv***) within the hypothalamus of *Mch-cre;L10-Egfp* mice. Representative sample of EGFP*^Mch^*-only cells (open arrowheads), EGFP*^Mch^* cells that coexpressed both CART and NK3R (EGFP*^Mch^*/CART/NK3R; filled arrowheads), EGFP*^Mch^* cells that coexpressed CART but not NK3R (EGFP*^Mch^*/CART; * asterisk), and EGFP*^Mch^*cells that were CART-negative but coexpressed NK3R (EGFP*^Mch^*/NK3R; † dagger). Percentage of EGFP*^Mch^*/CART (blue), EGFP*^Mch^*/CART/NK3R (orange), EGFP*^Mch^*/NK3R (purple), and EGFP*^Mch^*-only cells (turquoise) determined in female (***b***) and male (***c***) *Mch-cre;L10-Egfp* mice. Anteroposterior distribution of all EGFP*^Mch^* cell types at each level of the hypothalamus assigned in accordance with the *Allen Reference Atlas* (Dong, 2008; ***d***). Mediolateral distribution of all EGFP*^Mch^* cell types relative to the third ventricle (V3) in accordance with stereotaxic coordinate from the *Allen Reference Atlas* (***e***). Scale: 100 μm (***a***).

As CART-expressing MCH cells are expectedly more abundant in the medial hypothalamus (Broberger, 1999; Vrang et al., 1999; Elias et al., 2001; Croizier et al., 2010), we expressed the proportion of each subpopulation across the mediolateral axis of the hypothalamus in 0.5 mm increments away from the third ventricle. All defined EGFP*^Mch^* subtypes could be identified across the mediolateral axis of the hypothalamus, but overall, CART-positive EGFP*^Mch^* cells were more abundant medially then gradually diminishing laterally (**Figure 4e**). Interestingly, CART-positive cells that also expressed NK3R were evenly distributed mediolaterally (**Figure 4e**). By contrast, CART-negative EGFP*^Mch^*-only and EGFP*^Mch^*/NK3R cells gradually increased in abundance laterally (**Figure 4e**).

In order to determine the relative distribution of each EGFP*^Mch^* cell type, we used Nissl-based parcellations to delineate hypothalamic subnuclei and generate high-resolution spatial maps of all EGFP*^Mch^* cells. There were few EGFP*^Mch^*cells anteriorly (L63–L65), where they were loosely scattered medially and not confined to a specific hypothalamic nucleus (**Figure 5a, b**). The number of EGFP*^Mch^*/CART and EGFP*^Mch^*/CART/NK3R cells increased in L66–L69 and were laterally within the AHNp, the DMHa, and the ZI (**Figure 5c–f**). CART-negative EGFP*^Mch^*-only cells emerged and became more prominent posteriorly (L68–L73), where they were distributed lateral to the fornix in the LHA, in the region below the ZI and ventral border of the internal capsule and cerebral peduncle (**Figure 5e–j**). A few EGFP*^Mch^*/NK3R cells were scattered within this cluster of EGFP*^Mch^*-only cells (i.e., **Figure 5h**). Towards the posterior field (L72–74), EGFP*^Mch^* cells started to separate into two clusters. The medial cluster comprised more CART- positive EGFP*^Mch^*/CART and EGFP*^Mch^*/CART/NK3R cells, and they were distributed between the fornix and the DMH (**Figure 5i, j**) and along the ventral border of the posterior hypothalamic nucleus (PH; **Figure 5k**). A second cluster was lateral to the fornix and largely comprised EGFP*^Mch^*-only cells in the preparasubthalamic nucleus (PST; **Figure 5i**), ventral edge of the subthalamic nucleus (STN; **Figure 5i**), and parasubthalamic nucleus (PSTN; **Figure 5k**). The availability of EGFP*^Mch^* cells was sparse in posterior hypothalamic regions (L75–L77; **Figure 5l**).

**Figure 5.**
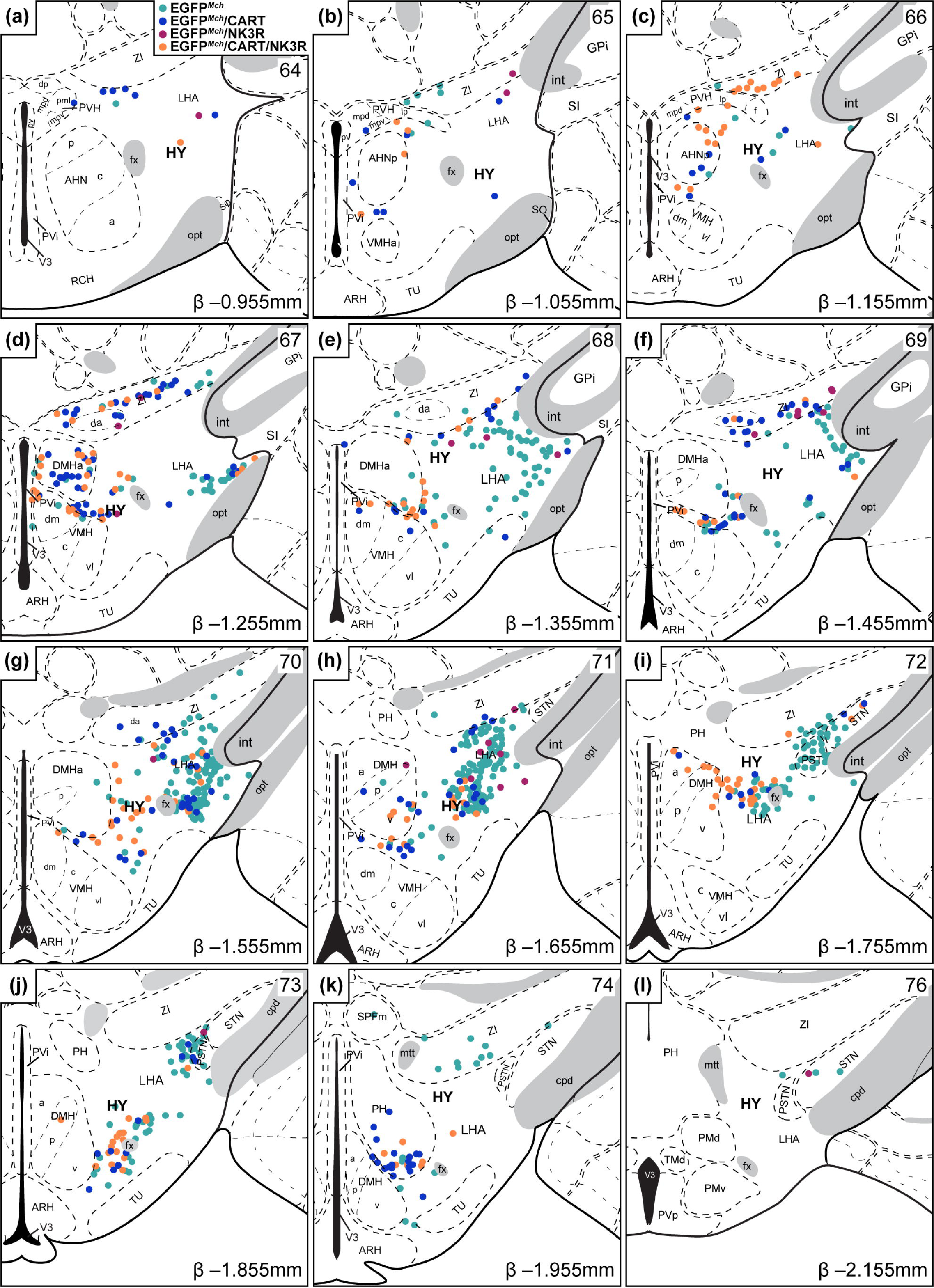
Spatial distribution of CART and/or NK3R coexpression in heterogeneous MCH cells. Representative coronal maps using *Allen Reference Atlas* (*ARA*; Dong, 2008) brain templates ordered from rostral to caudal (***a*–*l***) showing the relative distribution of EGFP*^Mch^*-only (turquoise circles), EGFP*^Mch^*/CART (blue circles), EGFP*^Mch^*/NK3R (purple circles), and EGFP*^Mch^*/CART/NK3R cells (orange circles) from the hypothalamus of *Mch-cre;L10-Egfp* mice. Each panel includes the corresponding *ARA* level (top right), stereotaxic coordinate inferred from Bregma (β; bottom right), and brain region labels according to *ARA* nomenclature.

As expected, our spatial maps revealed neuroanatomical division between medial CART- positive and lateral CART-negative MCH cells. However, there was no mediolateral separation for NK3R expression in CART-positive MCH cells. This topographical division suggested that the heterogeneity of MCH cells, especially based on CART coexpression, could contribute to unique neuronal networks.

## Female MCH cells differed in passive membrane properties

To determine if differences in electrophysiological characteristics, including passive and active membrane properties, also contribute to the overall heterogeneity of MCH cells, we performed whole-cell patch-clamp recordings from CART-positive (MCH/CART+) and CART-negative (MCH/CART−) MCH cells from *Mch-cre; L10-Egfp* male and female mice (**Figure 6a**). We recorded and biocytin-filled cells identified by EGFP*^Mch^* and processed them for post hoc immunostaining to determine the presence (**Figure 6b**) or absence of CART immunoreactivity (**Figure 6c**) in MCH/CART+ and MCH/CART− cells, respectively. We found prominent differences between MCH/CART+ and MCH/CART− cells based on their electrophysiological properties (**Supplemental Figure 1**), but the results herein were separated by sex to account for potential sex differences in the MCH system.

**Figure 6.**
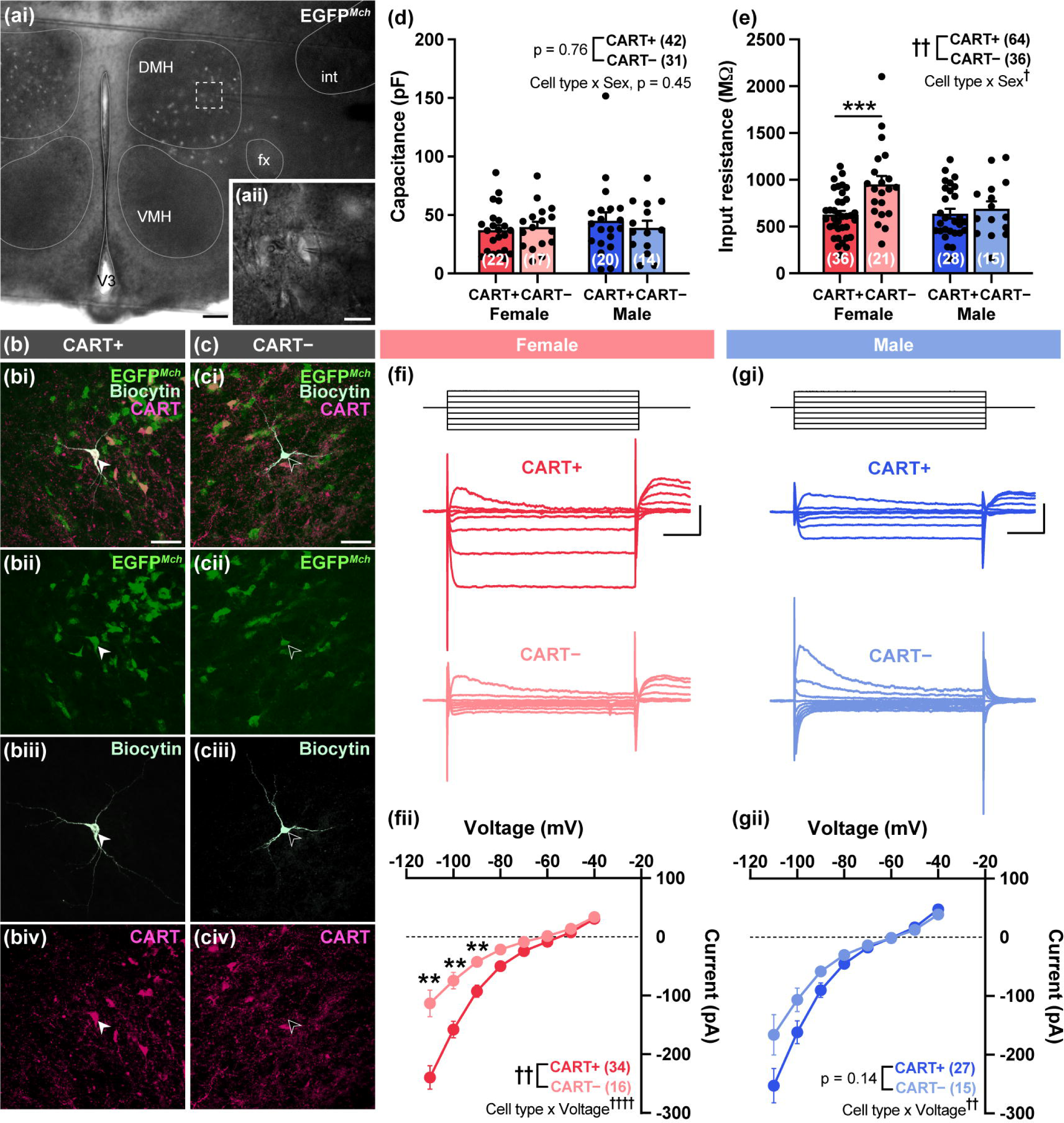
Lower input resistance and larger inward current elicited at female MCH/CART+ cells. Low (***ai***) and high magnification (***aii***, from outlined region in ***ai***) photomicrograph captured by infrared differential interference contrast imaging of a coronal brain slice from *Mch-cre;L10- Egfp* mice. Merged channel (***i***) confocal photomicrograph of native EGFP fluorescence (EGFP*^Mch^*; ***ii***), biocytin-labeling (***iii***), and CART immunoreactivity (***iv***) in MCH/CART+ (***b***; filled arrowhead) and MCH/CART− cell (***c***; open arrowhead). Comparison of capacitance (***d***), input resistance (***e***) and current–voltage relationship (***f*, *g***). Representative sample traces of voltage steps (*top*) and current elicited (*bottom*) in female (***fi***) and male (***gi***) MCH/CART+ and MCH/CART− cells. Current-voltage relationship elicited in female (***fii***) and male (***gii***) MCH/CART+ and MCH/CART− cells. Scale: 200 μm (***a***), 40 μm (***b****, **c***), 200 pA, 50 ms (***fi****, **gi***). Two-way ANOVA: †, p < 0.05; ††, p < 0.01; ††††, p < 0.0001 with Bonferroni multiple comparisons posttest: **, p < 0.01; ***, p < 0.001.

There was no difference in membrane capacitance of MCH cells by cell type (i.e., based on CART coexpression; F(1, 69) = 0.1, p = 0.759) or sex (F(1, 69) = 0.6, p = 0.452; **Figure 6d**). However, there was a strong effect of cell type on input resistance (F(1, 96) = 8.8, p = 0.004). The effect of sex on input resistance was less pronounced (F(1, 96) = 3.9, p = 0.051) but interacted with cell type to effect input resistance (F(1, 96) = 4.6, p = 0.034). We found that the mean input resistance of MCH/CART+ cells was lower than MCH/CART− cells in female (t = 3.9, df = 96, p = 0.003) but not male mice (t = 0.53, df = 96, p = 0.999; **Figure 6e**). Accordingly, the current–voltage relationship at female MCH cells differed by cell type (F(1, 47) = 12.0, p = 0.001) and interacted with voltage (F(7, 329) = 8.639,p < 0.0001), as a larger inward current was elicited at negative voltage potentials in MCH/CART+ cells (**Figure 6f**). In male MCH cells, there was no effect of cell type on current output (F(1, 39) = 2.2, p = 0.141), but cell type also interacted with voltage (F(7, 273) = 3.0, p = 0.004) as the current elicited tended to be greater at negative voltage steps (**Figure 6g**). To determine if there is a sex difference in the current– voltage relationship between MCH cells, we ran a three-way ANOVA with sex included as a factor with cell type and voltage. We found significant effects of voltage (F(1.110, 95.45) = 172.4, p < 0.0001) and cell type (F(1, 86) = 11.63, p = 0.001) on current output, but there was no overall effect of sex (F(1, 86) = 0.51, p = 0.477), and there was no significant interaction between sex and voltage (F(7, 602) = 1.2, p = 0.280) or cell type (F(1, 86) = 1.19, p = 0.279).

## MCH/CART+ cells exhibited pronounced spike rate adaptation

There was no effect of cell type (F(1, 110) = 0.2, p = 0.646) or sex (F(1, 110) = 1.2, p = 0.267) on the resting membrane potential of MCH cells, and there was also no interaction between cell type and sex (F(1, 110) = 0.2, p = 0.678). There were no resting membrane potential differences between the MCH/CART+ and MCH/CART− cells in female (t = 0.03, df = 62, p = 0.974) or male mice (t = 0.7, df =47.9, p = 0.490; **Figure 7a**).

**Figure 7.**
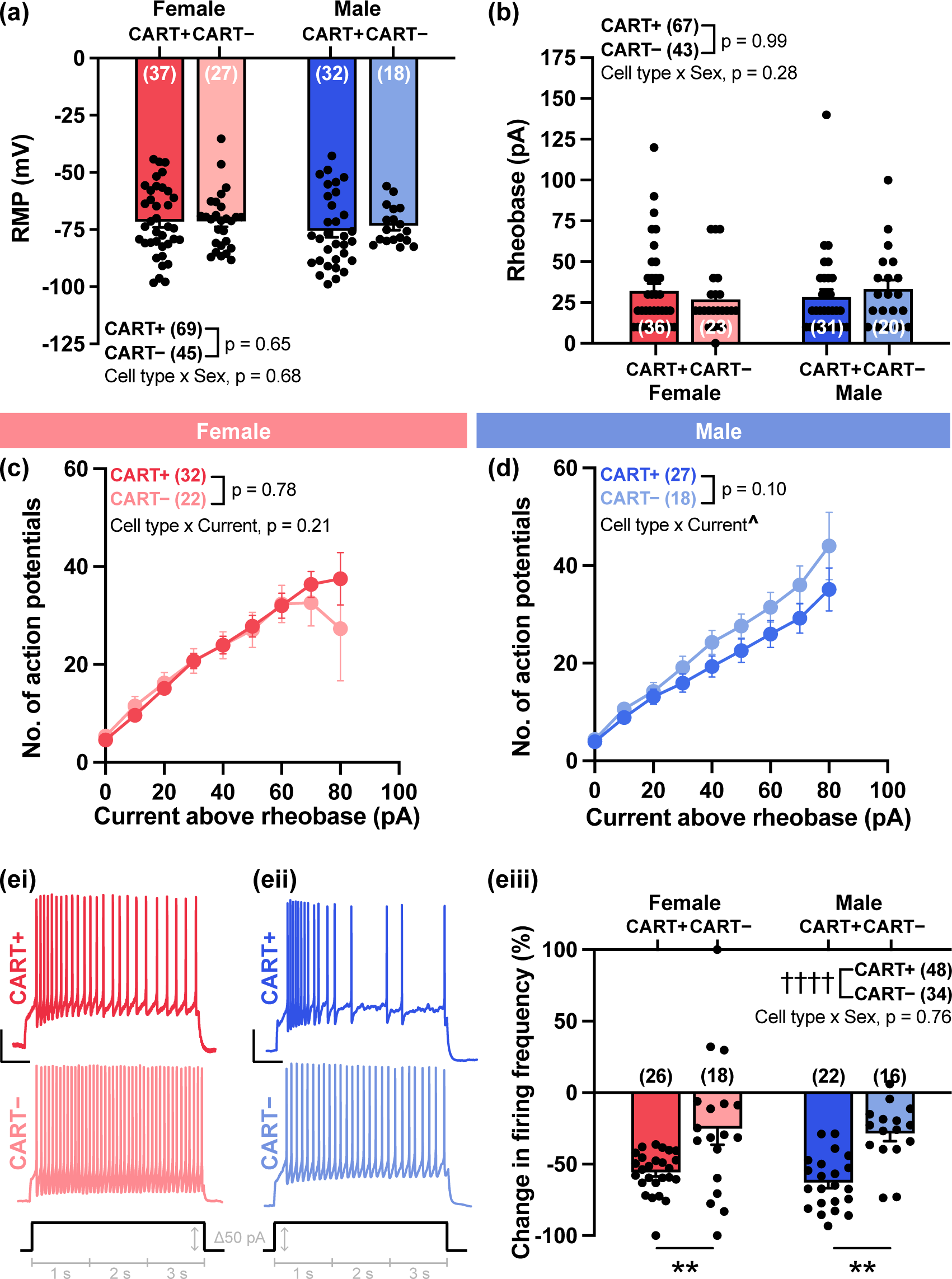
Elevated spike rate adaptation at MCH/CART+ cells. Comparison of resting membrane potential (RMP; ***a***), rheobase (***b***), and the number of action potentials elicited with increments of +10 pA current steps above rheobase between MCH/CART+ and MCH/CART− cells in female (***c***) and male *Mch-cre;L10-Egfp* mice (***d***). Representative sample current traces of action potential firing (***i***; *top*) elicited by a current step (3 s) at Δ50 pA above rheobase for its respective cell (*bottom*, black trace) from female (***ei***) and male cells (***eii***). Comparison of the percent change in instantaneous frequency of action potentials elicited at the start (first 1-s bin) and end (third 1-s bin) of the 3 s current step (***eiii***). Two-way ANOVA: ††††, p ≤ 0.0001 or mixed-effect ANOVA: ^, p < 0.05 with Bonferroni multiple comparisons posttest: **, p < 0.01. Scale: 20 mV, 500 ms (***ei*, *eii***).

While there were no differences in resting membrane potential, we determined if there were differences in the electrical fingerprint or excitability of MCH/CART+ and MCH/CART− cells. We first determined if there were differences in the rheobase of MCH cell types by comparing the minimum amount of current required to elicit action potential firing. We found no difference in the rheobase of MCH/CART+ and MCH/CART− cells in female (t = 0.8, df = 106, p = 0.846) or male mice (t = 0.7, df = 106, p = 0.938), as there was no effect of cell type (F(1, 106) = 0.0002, p = 0.987), sex (F(1, 106) = 0.1, p = 0.779), or cell type by sex interaction (F(1, 106) = 1.2, p = 0.283; **Figure 7b**).

All MCH cells increased the number of action potentials elicited with increasing current steps from the rheobase (female: F(8, 309) = 80.7, p < 0.0001; male: F(8, 274) = 123.1, p < 0.0001). Interestingly, while there was also no effect of cell type at female (F(1, 53) = 0.1, p = 0.783; **Figure 7c**) or male cells (F(1, 43) = 2.9, p = 0.097; **Figure 7d**), we found that cell firing depended on the amount of current injected at each cell type. There was a significant interaction between current injection and cell type at male (F(8, 274) = 2.5, p = 0.013) but not female MCH cells (F(8, 309) = 1.4, p = 0.213). To determine if there was a sex difference in the excitability of MCH cells, we included sex as a factor in an overall model with cell type and the amount of current injected. There was no main effect of sex (F(1, 95) = 0.4, p = 0.547), but sex was an important factor in a three-way interaction with cell type and current injection to impact action potential firing (F(8, 583) = 3.4, p = 0.001).

Finally, as the excitability of MCH cells may be higher in the juvenile period (Linehan and Hirasawa, 2018), we determined if the excitability of MCH cells were related to age. The change in firing frequency of female MCH/CART+ cells increased with age (*R^2^* = 0.22, p = 0.016) but no other active membrane properties were age-dependent (**Supplemental Table 1**).

As previously shown (Burdakov et al., 2005), all MCH cells exhibited spike rate adaptation. We compared the percent change in firing frequency at the start (first 1-s bin) and end (third 1-s bin) of the depolarizing current step to determine differences in firing (**Figure 7e*i*, e*ii***). We found a strong effect of cell type on the change in firing frequency (F(1, 78) = 28.1, p < 0.0001) but no effect of sex (F(1,78) = 0.7, p = 0.392; **Figure 7e*iii***) or interaction between sex and cell type (F(1, 78) = 0.1, p = 0.759). Interestingly, both male (t = 3.8, df = 78, p = 0.002) and female MCH/CART+ cells (t = 3.7, df = 78, p = 0.003) had a higher firing rate at the start than end of the current step (**Figure 7e*iii***). This was consistent with a unique electrical fingerprint in MCH/CART+ cells that comprised an initial burst of action potentials followed by a sharp drop in firing frequency (**Figure 7e**). Taken together, our results indicated that all MCH cells underwent spike rate adaptation upon sustained depolarization regardless of sex.

## Differences in rise time kinetics of excitatory events at MCH cells

The differences in spatial distribution and electrical properties of MCH/CART+ and MCH/CART− cells suggested that MCH cells may be differentially regulated by afferent input. In particular, excitatory inputs to MCH cells are regulated by feeding (Li and Van Den Pol, 2009; Linehan et al., 2020) and sleep (Briggs et al., 2018), thus we analyzed the spontaneous excitatory postsynaptic currents (sEPSC) arriving at MCH/CART+ and MCH/CART− cells in female (**Figure 8a**) and male brains (**Figure 8b**). Properties of sEPSC events did not vary with age (**Supplemental Table 1**).

**Figure 8.**
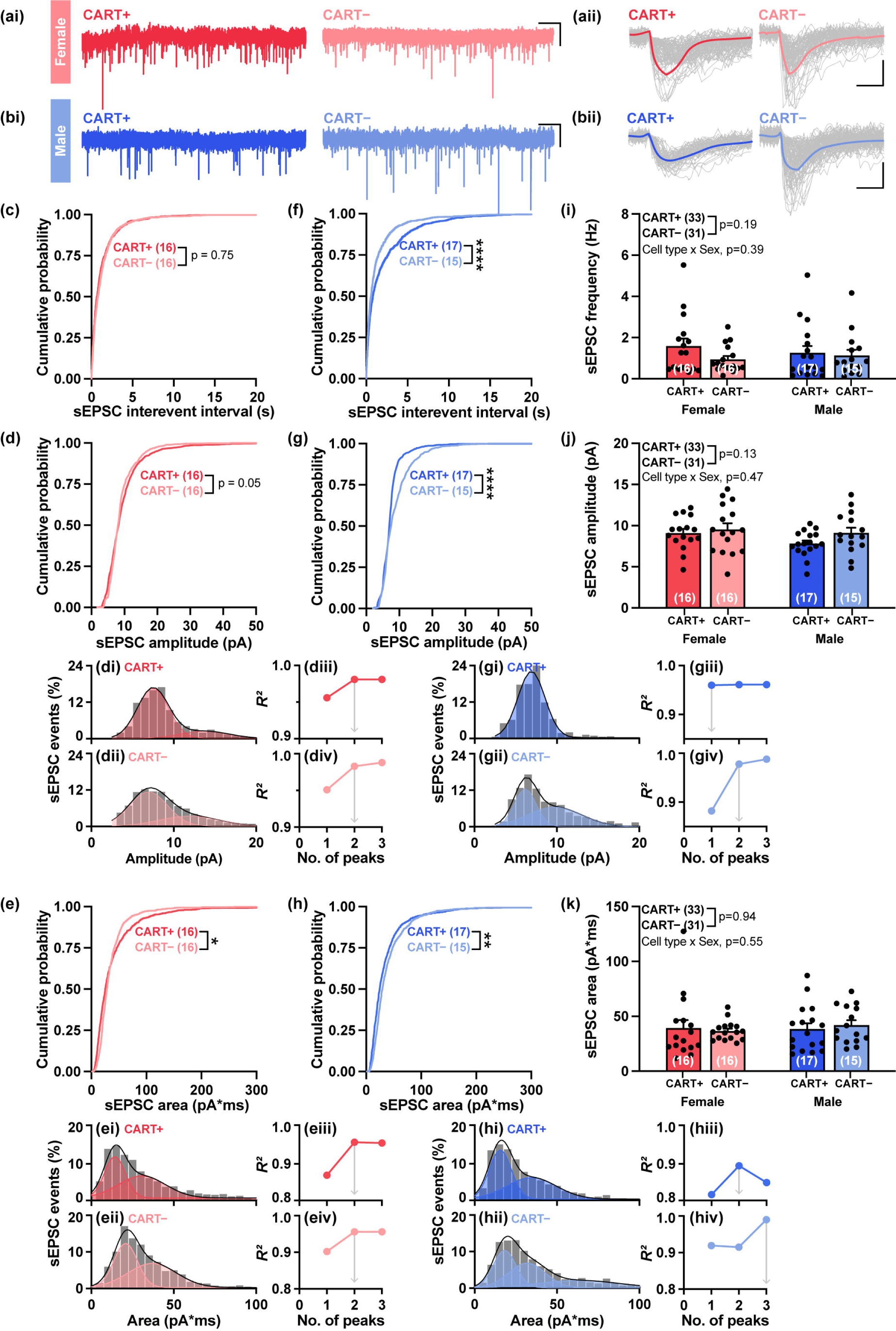
Smaller and less frequent sEPSC events at MCH/CART+ cells. Representative current trace (***i***) and sample of individual spontaneous excitatory postsynaptic current (sEPSC) events superimposed, with average trace shown in bold (***ii***), when recorded at a holding potential of V_h_ = −60 mV from female (***a***) and male (***b***) MCH/CART+ and CART− cells. Cumulative probability plot of the interevent intervals, amplitude, and area of sEPSC in female (***c*–*e***) and male cells (***f*–*h***). Overlay of the number of peaks (red and blue shaded regions) fitted against the normalized frequency distribution (gray bars) of sEPSC amplitudes (***d****, **g***) and sEPSC areas (***e****, **h***) at CART+ (***i***) and CART− (***ii***) cells with their corresponding goodness of fit (*R*^2^) at increasing number of curves fitted (***iii*, *iv***). Number of peaks that best fit to the sEPSC amplitude (***iii***) and area (***iv***) is indicated by the gray arrow. X-axis labels in ***i*** and ***iii*** correspond with that shown in ***ii*** and ***iv***, respectively. Mean of the frequency, amplitude, and area of sEPSC events (***i*–*k***). Scale: 5 s, 10 pA (***ai*, *bi***); 5 s, 5 pA (***aii*, *bii***).

We found no difference in the cumulative distribution of interevent intervals (p = 0.750; **Figure 8c**) or amplitude (p = 0.052; **Figure 8d**) of sEPSCs arriving at female MCH cells. However, we found a significant rightward shift of the cumulative probability of the area of sEPSC events (p = 0.032; **Figure 8e**) arriving at female MCH/CART+ cells. At male MCH cells, we found differences in the frequency and size of sEPSC events collected. There was a rightward shift in the cumulative probability of interevent intervals (p < 0.001; **Figure 8f**) and leftward shift in the cumulative probability of sEPSC amplitudes (p < 0.0001; **Figure 8g**) and areas (p = 0.007; **Figure 8h**) thus implicating that sEPSC events at male MCH/CART+ cells may arrive less frequently and be smaller. Indeed, a comparison of the normalized frequency distribution of sEPSC amplitudes (**Figure 8di, dii)** and areas (**Figure 8ei, eii**) in female cells revealed two underlying populations sampled for each cell type. In male CART+ cells, normalized frequency distributions displayed a single population of sEPSC amplitudes (**Figure 8gi**) and two populations of sEPSC areas (**Figure 8hi**). At male CART− cells, there was a greater variation in sEPSC amplitudes that were fit with two peaks (**Figure 8gii**) and sEPSC areas that were fit with three peaks (**Figure 8hii**). However, we found no significant effect of cell type on the mean sEPSC frequency (F(1, 60) = 1.8, p = 0.188; **Figure 8i**), amplitude (F(1, 60) = 2.3, p = 0.133; **Figure 8j**), or area (F(1, 60) = 0.01, p = 0.937; **Figure 8k**). There was also no sex difference on the mean sEPSC frequency (F(1, 60) = 0.1, p = 0.801; **Figure 8i**), amplitude (F(1, 60) = 2.3, p = 0.138; **Figure 8j**), or area (F(1, 60) = 0.2, p = 0.629; **Figure 8k**), and no significant interactions between cell type and sex on these sEPSC parameters.

Interestingly, we found differences in sEPSC kinetics at female and male MCH cells, as a rightward shift in the cumulative distribution of sEPSC rise time rate (i.e., amount of time it takes for a sEPSC event to cover 10–90% of its amplitude; **Figure 9a**) at female (p < 0.0001; **Figure 9b**) and male MCH/CART+ cells (p < 0.0001; **Figure 9c**) indicated that a subset of sEPSC events at MCH/CART+ cells had slower rise rate than MCH/CART− cells. Cell type (F(1, 55) = 7.0, p = 0.010) and sex (F(1, 55) = 4.9, p = 0.030) were significant factors influencing rise time kinetics but there was no interaction to influence the rising rate of sEPSC events (F(1, 55) = 0.3, p = 0.578). In effect, the mean sEPSC rise rate at MCH/CART+ cells in female (t = 1.7, df = 26.18, p = 0.104) and male mice (t = 2.0, df = 24.06, p = 0.052) tend to be slower than at MCH/CART− cells (**Figure 9d**).

**Figure 9.**
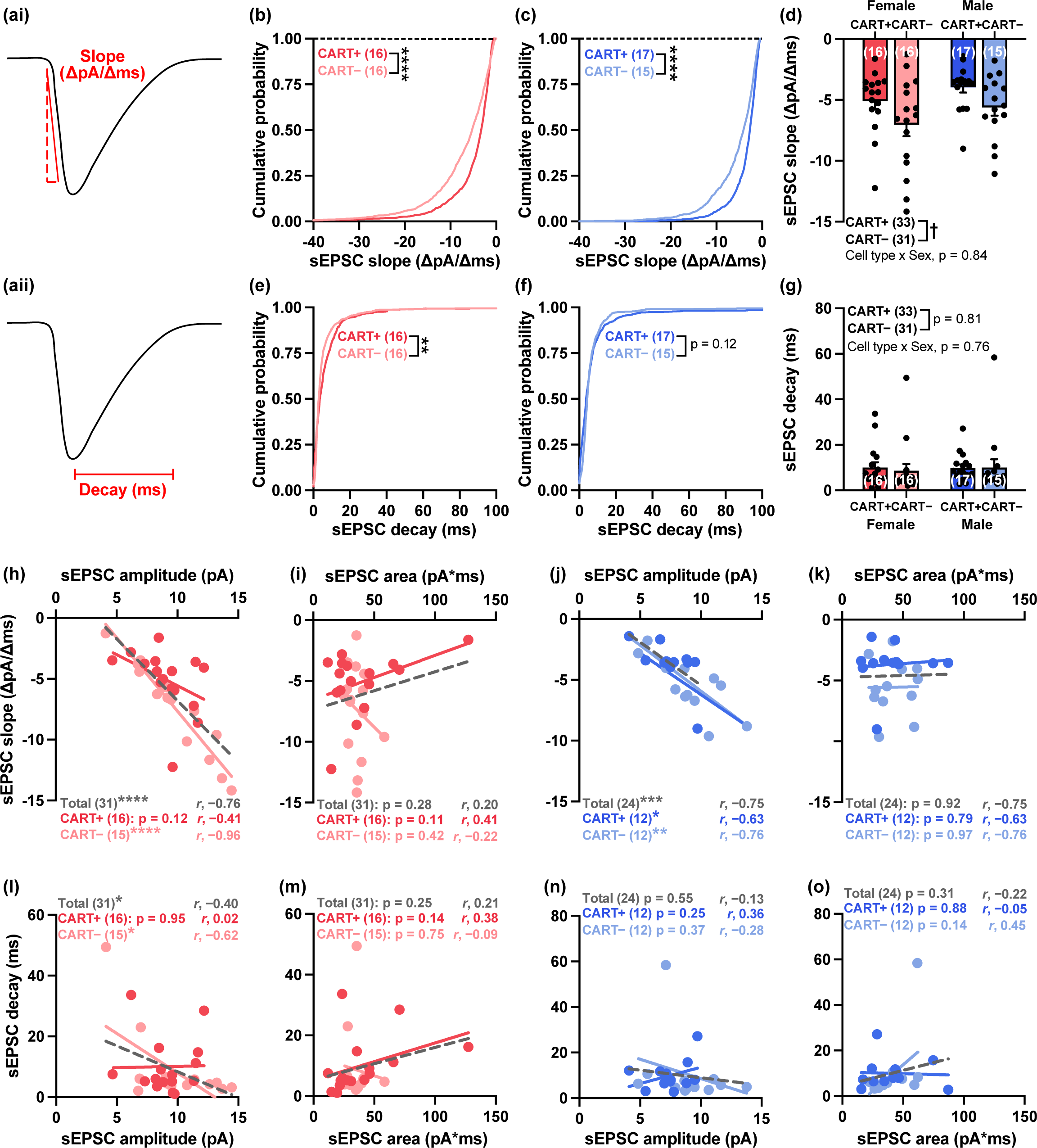
Slower sEPSC event kinetics at MCH/CART+ cells. Drawing of a sEPSC event illustrating the analysis of rise slope, measured as the change in amplitude over 10–90% of peak amplitude (***ai***), and decay time (***aii***). Cumulative probability plot and mean of sEPSC event slope (***b*–*d***) and decay (***e*–*g***) in female and male cells. Correlation plots of the relationship between sEPSC slope versus amplitude (***h*, *j***) or area (***i*, *k***) and between sEPSC decay versus amplitude (***l*, *n***) or area (***m*, *o***) for CART+ (red, dark blue), CART− (pink, light blue) or all MCH cells (grey). Two-way ANOVA: †, p < 0.05. Kolmogorov-Smirnov test: **, p < 0.01; ****, p < 0.0001. Lines representing Pearson correlation are denoted by *r*, with significance determined by simple linear regression: *, p < 0.05; **, p < 0.01; ***, p < 0.001; ****, p < 0.0001.

By contrast, differences in decay time of sEPSC events were less prominent. There was a slight rightward shift in the cumulative distribution of sEPSC decay times of at female MCH/CART+ cells (p = 0.001; **Figure 9e**), thus suggesting that a small subset of sEPSC events had longer or slower decay times. No differences were observed between male MCH cells (p = 0.122; **Figure 9f**). However, there was no effect of cell type (F(1, 55) = 0.1, p = 0.802), sex (F(1, 55) = 0.04, p = 0.833), or interactions between cell type and sex (F(1, 55) = 0.1, p = 0.797; **Figure 9g**).

The rise kinetics of sEPSC events were directly correlated with their amplitude in most MCH cells. Slow rising sEPSC events tended to be smaller in amplitude at female (**Figure 9h**) and male MCH cells (**Figure 9j**) but was not related to sEPSC area (**Figure 9i, 9k**). The sEPSC decay times were largely unrelated to sEPSC amplitude (**Figure 9l, 9n**) or area (**Figure 9m, o**). As these relationships were seen at most MCH cell types, it is unlikely that any differences would be related to CART expression.

Overall, our findings indicated that there were no gross differences in the frequency, amplitude, or area of sEPSC events at either subtype or sex of MCH cells, though a subset of events at male CART+ cells may occur less frequently and have smaller amplitudes. Interestingly, we found cell type-dependent differences in the kinetics of sEPSC events, as those arriving at CART+ cells had slower rise rates in both sexes, and a subset of events may also have slower decay times. Similarly, when sEPSC events from each sex were pooled, we also did not find prominent differences between MCH/CART+ and MCH/CART− cells based on the properties of their afferent input (**Supplemental Figure 2**).

## Male MCH/CART– were highly branched

To determine if the properties of sEPSC events were a function of cell morphology, we first determined CART immunoreactivity in biocytin-filled EGFP*^Mch^* cells (**Figure 10a, b**) from whole-cell recordings (**Figure 6a–c**). Next, we used nickel-enhanced DAB staining to label the soma, axons, and dendrites of each cell (**Figure 10c**) so they may be reconstructed to evaluate dendritic branching patterns using the Sholl analysis (**Figure 10d–f**). The branched structure of MCH cells was not related to membrane capacitance (**Supplemental Figure 3**). The dendritic features of MCH cells were comparable between CART+ and CART− cells (**Supplemental Figure 4**), but datasets from male and female mice were separated to assess if there were sex-specific differences between CART+ and CART− cells.

**Figure 10.**
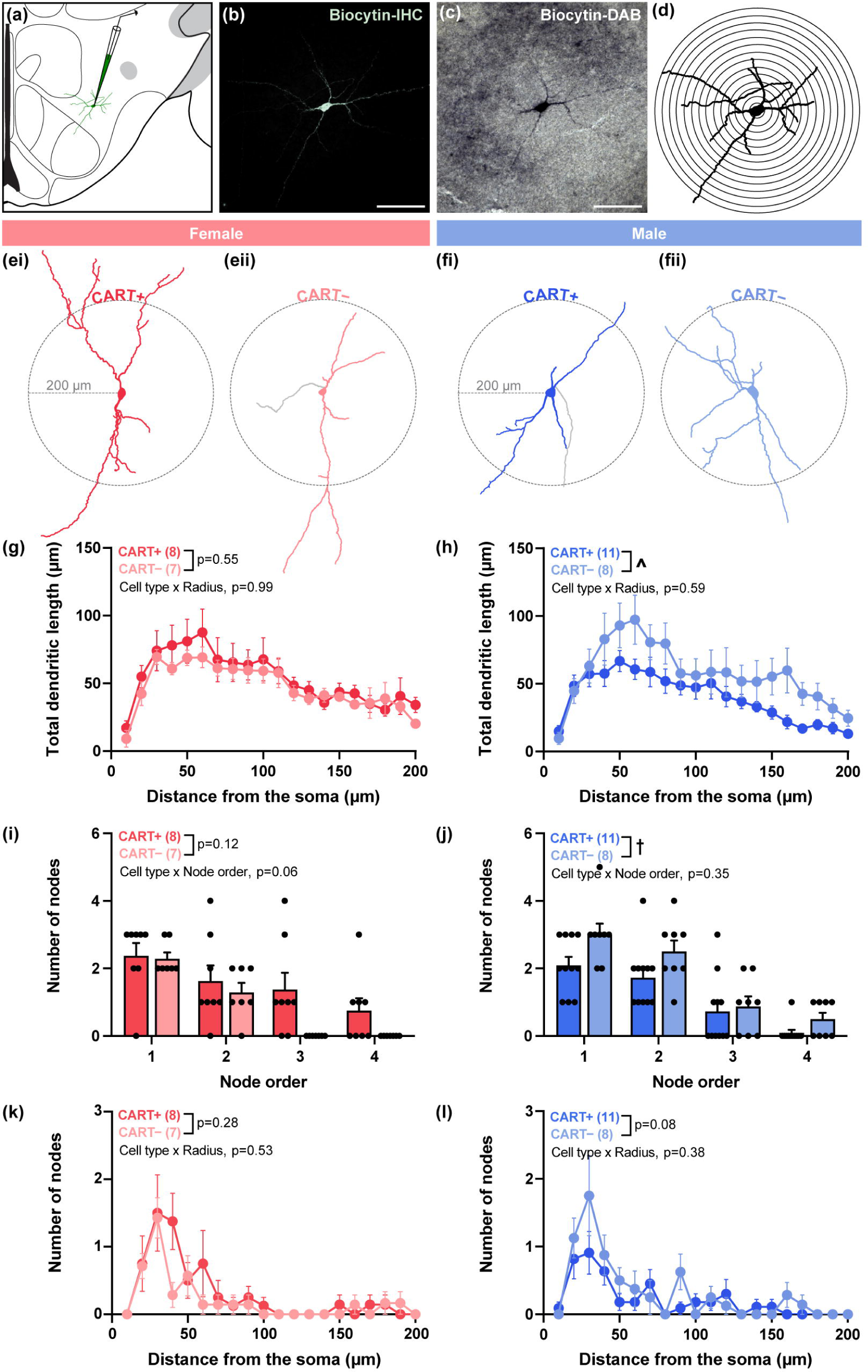
Lower branching and dendritic coverage at male MCH/CART+ cells. Schematic of a whole-cell recording at an EGFP*^Mch^* cell in *Mch-cre;L10-Egfp* brain tissue that was biocytin-filled during the recording (***a***). Biocytin was first labelled in fluorescence (***b***) to determine the presence or absence of CART coexpression (not shown) and then nickel-enhanced with DAB staining (***c***) to trace and reconstruct the cell for Sholl analysis (***d***). Representative reconstructed female (***e***) and male (***f***) MCH/CART+ (***i***) and MCH/CART− (***ii***) cells with axons shown in gray. Dashed circles outline the field analyzed within a 200 μm radius from the soma. Comparison of total dendritic length coverage (***g*, *h***), number of nodes per node order (***i*, *j***), or distance of nodes from the soma (***k*, *l***). Two-way ANOVA: †, p < 0.05 or mixed-effect ANOVA: ^, p < 0.05.

Cell type had no effect on the total dendritic length of female MCH cells (F(1, 13) = 0.4, p = 0.545; **Figure 10g**) but influenced that of male MCH cells (F(1, 17) = 6.9, p = 0.018; **Figure 10h**). To determine if there was an overall sex difference, we included sex in our model with cell type and distance, but we did not see an effect of sex (F(1, 28) = 0.04, p = 0.845), cell type (F(1, 28) = 0.1, p = 0.816), or any interactions involving sex or cell type on dendritic length.

We also used cell reconstructions to determine the number of nodes (i.e., branch points) at each node order (e.g., primary, secondary, tertiary, quaternary branch). Female cells did not show an effect of cell type on the number of nodes at each increasing node order (F(1, 13) = 2.7, p = 0.123; **Figure 10i**), but we found an effect of cell type at male cells (F(1, 17) = 5.0, p = 0.040), as male MCH/CART+ cells tended to have fewer nodes at each branch order (**Figure 10j**). However, there was no overall sex difference in the number of nodes (F(1, 30) = 1.0, p = 0.316), but in a three-way ANOVA that included sex, cell type, and node order, we found that CART expression significantly interacted with sex (F(1, 30) = 7.3, p = 0.011) and also with node order (F(3, 90) = 3.6, p = 0.016). Therefore, while male CART+ cells tended to have less nodes at each node order compared to CART− cells, female cells may show the opposite or no pattern.

Given that there may be a difference in the number of branch points in male cells, we also determined if the number of nodes varied as a function of distance from the soma. There were no differences in the number of nodes at any distance away between CART+ and CART− female (F(1, 13) = 1.2, p = 0.284; **Figure 10k**) or male cells (F(1, 312) = 3.1, p = 0.080; **Figure 10l**). There was no sex difference (F(1, 30) = 0.05, p = 0.629), but the dendrites of MCH cells can have different node distributions depending on sex (F(1, 28) = 4.4, p = 0.044).

Taken together, as male MCH/CART+ cells had less dendritic length and fewer branch points, they may have a less complex morphological structure. Meanwhile, female MCH cells did not appear to be morphologically distinct from each other.

## CART-dependent association between synaptic events and dendritic structure

As dendritic structure can impact synaptic activity and vice versa (Spruston et al., 2016), we next determined if the properties of sEPSC events, including frequency (**Figure 11a–d**), amplitude (**Figure 11e–h**), area (**Figure 11i–l**), rise slope (**Figure 11m–p**), and decay (**Figure 11q–t**), were related to the dendritic morphology of CART+ or CART− MCH cells.

**Figure 11.**
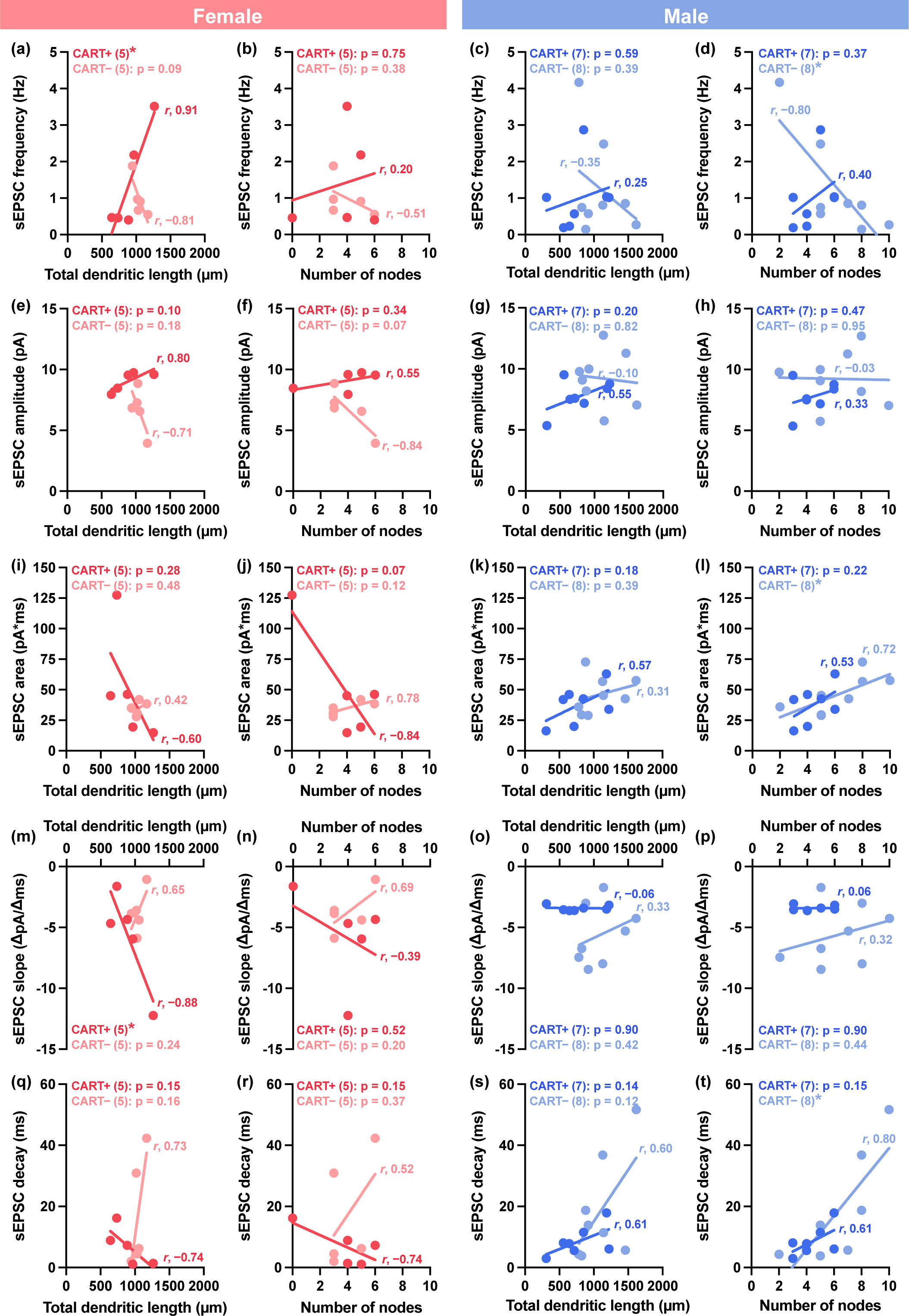
Dendritic branching predicted lower frequency sEPSC events with larger area at male MCH/CART− cells. Correlational plots of sEPSC frequency (***a****–**d***), amplitude (***e****–**h***), area (***i****–**l***), rise slope (***m****–**p***), and decay time (***q****–**t***) with dendritic length and number of nodes in MCH/CART+ or MCH/CART− cells. Lines representing Pearson correlations are denoted by *r*, with significance determined by simple linear regression: *, p < 0.05.

MCH/CART+ cells were best distinguished in female mice, as greater total dendritic length was correlated with the higher frequency of sEPSC events (*R*^2^ = 0.84, p = 0.030; **Figure 11a**) and faster rise kinetics (*R*^2^ = 0.78, p = 0.046; **Figure 11m**). Dendritic length was not correlated with sEPSC amplitude (*R*^2^ = 0.65, p = 0.101; **Figure 11e**), area (*R*^2^ = 0.37, p = 0.280; **Figure 11i**), or decay kinetics (*R*^2^ = 0.55, p = 0.149; **Figure 11q**). By contrast, none of the sEPSC parameters were related to dendritic length at female MCH/CART− cells (**Figure 11a, e, i, m, q**). sEPSC event properties were not related to dendritic nodes at neither female CART+ nor CART− cells (**Figure 11b, f, j, n, r**).

MCH/CART− cells were most distinct in male mice. There were no significant relationships between sEPSC properties and total dendritic length at CART+ or CART− cells (**Figure 11c, g, k, o, s**). However, in CART– cells we found that the number of dendritic nodes was inversely related with sEPSC frequency (*R*^2^ = 0.63, p = 0.018; **Figure 11d**) but directly related with area (*R*^2^ = 0.52, p = 0.045; **Figure 11l**) and decay time (*R*^2^ = 0.64, p = 0.018; **Figure 11t**). Meanwhile, no significant relationships could be discerned between sEPSC events and dendritic nodes at male CART+ cells.

In sum, the dendritic length was positively correlated with the sEPSCs frequency and slope at female MCH/CART+ cells, while the number of dendritic nodes was significantly related to the frequency, area, and decay kinetics of sEPSC events arriving at male MCH/CART− cells.

### DISCUSSION

This study aimed to characterize the neuroanatomical distribution, passive and active electrophysiological properties, and dendritic morphology of MCH/CART+ and MCH/CART− cells in male and female mice. There were no gross sex differences in the distribution or properties of MCH cells, however sex interacted with CART coexpression to influence the electrical fingerprint and dendritic branching of MCH cells. Furthermore, MCH/CART+ and MCH/CART− cells also differed based on their cellular conductance and on the kinetic properties of incoming synaptic events. Our findings suggested that each MCH cell type could form distinct neural circuits and elicit functionally distinct roles in the expression of MCH- mediated behaviours.

## Neuroanatomy: Mapping CART and NK3R expression in hypothalamic MCH cells

About half of the hypothalamic MCH cell population expressed CART. The prominence of CART coexpression in half of MCH cells has also been independently reported by others (Broberger, 1999; Elias et al., 2001; Cvetkovic et al., 2004; Vrang, 2006; Croizier et al., 2010; Mickelsen et al., 2017) while *Cartpt* mRNA coexpression may be even higher (Mickelsen et al., 2019; Fujita et al., 2021). CART-positive MCH cells can also be spatially differentiated from CART-negative cells. Consistent with previous reports (Broberger, 1999; Vrang et al., 1999; Brischoux et al., 2001; Wang et al., 2021), CART-positive MCH cells appeared more anteriorly and were largely distributed medial to the fornix within the anterior DMH, the medial ZI, and medial regions of the LHA. Meanwhile, CART-negative MCH cells were predominantly located in the lateral ZI and lateral regions of the LHA, near the internal capsule, the substantia innominata, and the cerebral peduncle.

Interestingly, NK3R expression defined another subset of MCH cells, as NK3R was almost exclusively seen in MCH/CART+ cells, and only a few MCH cells expressed NK3R in the absence of CART. This is consistent with single-cell RNAseq analysis (Mickelsen et al., 2019; Fujita et al., 2021) revealing that *Pmch* cells can be divided into two major subclusters: one that strongly expressed both *Carpt* and *Tacr3*, the genes for CART and NK3R respectively, and one that was *Cartpt*- and *Tacr3*-negative. Furthermore, *Cartpt* and *Tacr3* have also been recognized as highly expressed genes in *Pmch* cells by other RNAseq datasets like HypoMap (Steuernagel et al., 2022). CART and NK3R protein coexpression in MCH cells has been demonstrated (Broberger, 1999; Vrang et al., 1999; Brischoux et al., 2001, 2002; Elias et al., 2001; Cvetkovic et al., 2004; Croizier et al., 2012; Wang et al., 2021) and might be even more prominent than the coexpression of *Cartpt* and *Tacr3*, which can comprise over 70% of MCH cells (Mickelsen et al., 2019). Interestingly, MCH/CART/NK3R+ cells were equally distributed along the medio-lateral axis of the hypothalamus. However, these NK3R-expressing MCH cells may mediate different functions depending on their spatial distribution because the medial and lateral regions of the hypothalamus can receive unique neurokinin B (NKB) projections (Cvetkovic et al., 2003). The medial hypothalamus was mostly innervated by NKB axons from the lateral septal complex, multiple hypothalamic nuclei, and periaqueductal grey. In contrast, the lateral regions received the densest NKB innervation from the lateral hypothalamus and the pallidum, especially the substantia innominata and the diagonal band nucleus. In addition, Fujita et al. (2021) also recently identified the bed nucleus of the stria terminalis and central amygdala as major sources of NKB innervation, and these projections were densest at the lateral border of the hypothalamus. Therefore, while MCH/CART/NK3R+ cells appear equally distributed along the mediolateral axis, they may be differentially innervated.

As CART-positive and CART-negative MCH cells had distinct spatial distributions, they may also have different efferent projection targets and recruit unique behavioural networks. Specific tracing experiments have highlighted targets of CART-positive MCH cells within the mesolimbic pathway, including the accumbens (Ekstrand et al., 2014) and ventral tegmental area (Dallvechia-Adams et al., 2002; Philpot et al., 2005), where half of MCH-ir varicosities colocalized with CART (Dallvechia-Adams et al., 2002). Both MCH and CART can modulate energy balance through the mesolimbic pathway, but these peptides have opposing effects. MCH injection into the accumbens shell of rats increased chow consumption (Georgescu et al., 2005), but similar injections of CART peptides decreased feeding (Yang et al., 2005). Consistently, co-injection of MCH and CART in the accumbens prevented CART-induced dopamine release in this region (Yang and Shieh, 2005), thus it is possible that coincident MCH and CART release in the accumbens could self-regulate to modulate feeding outcomes (Diniz and Bittencourt, 2017).

## Electrophysiology: Electrical fingerprint of MCH cells varied with sex

There were no overt sex differences in the passive or active membrane properties between MCH cell types, but differences between MCH cell types were apparent depending on the sex of the animal. The input resistance was lower in CART-positive MCH cells from female mice, so they may require a stronger current input to be stimulated or the same current input would elicit a lower voltage response in these female cells. MCH cells receive direct inputs from diverse brain regions at varying intensities (González et al., 2016). Thus, differences in input resistance may complement the strength of afferent inputs that innervate MCH cells so that, for instance, weak inputs at CART-negative MCH cells that have higher input resistances may be sufficient for functional activation. Notably, MCH cells largely differed based on their capacity for excitation. CART-positive MCH cells exhibited burst firing that dissipated with sustained depolarization, and this was seen by their pronounced spike rate adaptation.

Our findings extended those by Fujita and colleagues (2021) who noted that *Pmch* cells may express at least two electrical fingerprints, but these two fingerprints were not exclusively correlated to *Carpt* and *Tacr3* coexpression. Interestingly, we noted that *Tacr3* expression may be higher than the expression of its corresponding protein NK3R. As such, it is possible that electrical subclustering of *Pmch* cells better correlates with protein expression or more broadly based on CART expression. We found a greater proportion of MCH/CART+ cells displaying burst firing. The firing pattern of a neuron can influence both the target postsynaptic cell and the originating presynaptic cell. For example, different stimulation frequencies can influence gene expression in the target cell (Klein et al., 2003; Lee et al., 2017; Iacobas et al., 2019). At the presynaptic terminal, select firing patterns may differentiate the release of coexpressed chemical messengers. MCH neurons can utilize fast-acting neurotransmitters (Jego et al., 2013; Chee et al., 2015; Mickelsen et al., 2019; Sankhe et al., 2022), but they also coexpress multiple neuropeptides. Bursts of action potentials more effectively stimulate neuropeptide release than chronic firing (Poulain and Wakerley, 1982), and the amount of neuropeptide released per action potential is directly related to the firing frequency (Dreifuss et al., 1972; Gainer et al., 1986). Therefore, we predict that cell firing is imperative for their function. As both short, high frequency stimulation (Jiang et al., 2020) and sustained depolarization may be required for MCH release (Hausen et al., 2016; Noble et al., 2018), burst firing at CART-positive MCH cells may initiate neuropeptide release that is sustained with high frequency firing to permit maximal neuropeptide signaling (Jiang et al., 2020).

## Morphology: Differential dendritic structure at MCH cells varied with sex

There is little data on the dendritic branching pattern of MCH cells, which have been reported to be large and multipolar in hypothalamic cultures (Eggermann et al., 2003). Differences in the morphology between CART-positive and CART-negative MCH cells were most prominent in males, where CART-positive cells had fewer branch points, and these dendritic nodes were farther from the soma. In effect, we can infer that male CART-positive cells would have a smaller dendritic tree. Computer modelling studies have reported that the morphology of a neuron can directly impact the generation of outputs. In cortical and hippocampal neuron models, firing phenotype could be predicted by dendritic branching, where larger dendritic trees were less likely to display burst firing (Krichmar et al., 2002; Van Elburg and Van Ooyen, 2010; Psarrou et al., 2014). Consistent with these cortical models, our male CART-negative MCH cells, which had larger dendritic arborizations, were also less likely to exhibit burst firing when depolarized.

Estrogen and glutamate are important modulators of dendritic structure. Male cells in the ventromedial hypothalamic nucleus are more branched than female cells, and this effect was due to an estradiol-mediated increase in glutamatergic signalling (Mong and McCarthy, 1999; Shwarz and McCarthy, 2008). MCH cells are sensitive to the effects of chronic, but not acute (Tritos et al., 2004), estrogen treatment that blocked fasting-induced increases in *Pmch* gene expression (Murray et al., 2000; Mystkowski et al., 2000; Morton et al., 2004). As CART- negative MCH cells in the male brain were more branched, it suggested that they may be more sensitive to the influence of estrogen. Furthermore, as differences in branching were most prominent in the male brain, it is also possible that these differences were hardwired by estradiol during the sexual differentiation and masculinization periods of the developing brain (McCarthy, 2008).

Glutamatergic signalling is a key factor in dendritic development (Rajan and Cline, 1998; Sin et al., 2002; Richards et al., 2005). There were no significant differences in the average frequency or size of excitatory events to MCH cells in both sexes. However, we did detect a sample of glutamatergic events in male CART-negative cells that occur at a higher frequency with greater amplitude and area, and we can speculate that this could promote increased dendritic growth in these cells. MCH cells receive higher glutamatergic input during development (Li and Van Den Pol, 2009) or in the juvenile period (Linehan and Hirasawa, 2018), following sleep deprivation (Briggs et al., 2018), or high fat feeding (Linehan et al., 2020). Furthermore, hypothalamic cells can also increase dendritic branching following restraint stress (Grafe et al., 2019), thus behavioural experiences may shape the branching pattern and contribute to the plasticity of MCH cells even in adulthood.

## Intersection of form and function in MCH cells

The structure of a neuron can have a substantial impact on its electrical behaviour and help define the role it plays in a larger synaptic network. One discernible outcome at MCH/CART+ cells, especially in female mice, is that greater dendritic length was associated with higher sEPSC frequency but slower sEPSC kinetics. Greater dendritic lengths would reflect a larger surface available for synapses, but sEPSC events that synapse farther away from the soma would arrive with slower rise and decay times.

Conversely, dendrites may also converge at nodes, which act as points of synaptic integration, thus two synaptic events arriving from separate dendritic branches can converge and summate at a node to elicit synergistic effects and greater potential change than their independent events (Kamijo et al., 2014). This was notable at CART− cells and best shown in male mice where dendritic branching positively correlated with sEPSC area but negatively correlated with sEPSC frequency. The summation of synaptic events may reflect the direct positive relationship between sEPSC area and the availability of nodes at MCH/CART− cells that received a higher proportion of larger amplitude sEPSC events. Furthermore, the prevalence of sEPSC event summation may result in a lower frequency of sEPSC events detected at highly branched cells. By contrast, nodes can also be points of synaptic attenuation, especially as synaptic events travel from smaller branches with high intracellular resistance to larger diameter branches of proximal dendrites where current can be attenuated by charging larger membrane capacitances (Spruston et al., 2016). Therefore, synaptic events at distal dendrites of MCH/CART− cells that are highly branched may exhibit increased decay times as current dissipates along the dendrite. Paradoxically, while cells with greater dendritic branching may present more synaptic sites, there was an inverse relationship between sEPSC events frequency and dendritic nodes at MCH/CART− cells. This may be attributed to the attenuation of sEPSC events arriving at distal dendrites of highly branched cells and to the summation of smaller sEPSCs into larger events at dendritic nodes.

## Technical considerations

Majority of MCH cells (> 90%) could be identified by EGFP*^Mch^* in the hypothalamus of *Mch-cre;L10-Egfp* mice, but as previously shown (Beekly et al., 2020), some EGFP*^Mch^*cells, though infrequent and sporadically distributed, did not express MCH immunoreactivity. There was a unique perifornical cluster of MCH-negative EGFP*^Mch^*cells that were often grouped in a circular pattern immediately dorsal to the fornix at *ARA* L73. Our work expanded upon the initial analysis by Beekly and colleagues by defining the level that these MCH-negative EGFP*^Mch^* cells occur relative to Bregma. Interestingly, we found that this MCH-negative cluster also did not express *Pmch* or *Egfp* mRNA, so they were not actively producing EGFP. As EGFP production in the *Mch-cre;L10-Egfp* mouse arose from a fate-mapping strategy to mark the genetic lineage of a cell (Zinyk et al., 1998; Padilla et al., 2010), EGFP fluorescence would permanently mark cells that expressed *Pmch* at any point during gestation or development, even if *Pmch* transcription has since been turned off in the cell. EGFP fluorescence in MCH-negative cells may reflect legacy expression of *Mch* promoter activity, such as during development. We did not record from EGFP-labelled cells immediately surrounding the fornix, but it is possible that a MCH- negative EGFP-labeled cell may have been inadvertently included in our dataset.

In order to draw stronger associations between electrophysiological and morphological properties, we reconstructed the dendritic structure of MCH cells that we recorded from. The addition of dendritic spine analysis would help integrate and interpret differences in excitatory input between MCH cell types. However, we were unable to analyze the type or density of dendritic spines on our reconstructed cells because spines were not always visible with biocytin-labelling. Our reconstructed cells underwent extensive post hoc thick-tissue processing that may be prohibitive for maintaining the integrity of dendritic spines. However, sex-dependent differences in spine properties within other hypothalamic nuclei can relay functional outcomes on male and female behaviour (Matsumoto and Arai, 1980; Mong and McCarthy, 1999; Amateau and McCarthy, 2002, 2004) and would be relevant in future studies to further elucidate the heterogeneity among MCH cells. Furthermore, distal dendritic processes severed during dissection and tissue processing may underreport differences at distal dendrites. However, to account for this potential variation distally, we limited our morphological analyses to a 200-μm radius from the soma. Importantly, our results thus far indicated that the most dynamic differences in the dendritic tree occurred proximally within 50 μm of the soma.

The excitability of rat MCH cells may decrease after the juvenile period (> 7 weeks of age) (Linehan and Hirasawa, 2018). We recorded from a wide age range of animals, but as only 3% (2/69) of our male cells and 21% (12/57) of female cells were from mice under 7 weeks old, it was difficult for us to assess changes in excitability from the juvenile period. However, overall, neither the excitability of MCH cells or excitatory input to MCH cells varied with age in our dataset (**Supplemental Table 1**).

### CONCLUSION

The coexpression of CART within a subset of MCH cells had previously been established (Broberger, 1999; Elias et al., 2001; Cvetkovic et al., 2004; Vrang, 2006; Croizier et al., 2010; Mickelsen et al., 2017), and *Carpt/Tacr3* expression defines a chemically-distinct subpopulation of *Pmch* cells (Mickelsen et al., 2019). Thus, the main objective of this study was to assess potential differences between CART+ and CART– MCH cells. Overall, although there was no effect of sex, we conclude that these cells are phenotypically different, as we observed a significant effect of cell type on conductance, excitability, and sEPSC kinetics. However, it was important to assess for potential sex differences because the MCH system is known to be sexually dimorphic across several behaviours. A unique profile can be identified for each MCH cell type. CART-positive cells coexpressed NK3R, had lower input resistance, and greater spike rate adaptation. Meanwhile, CART-negative cells had higher cell resistance and displayed steady firing.

Differences in electrophysiological and morphological outcomes help elucidate the neural circuit involving CART-positive or CART-negative MCH cells. For instance, CART-positive cells may be innervated by strong afferent input and encode behaviours with short bursts of high frequency stimulation. By contrast, CART-negative cells may be sensitive to even weak or subthreshold afferent input but provide sustained cell firing to mediate long-lasting behavioural output. Interestingly, in males, CART-negative cells have more complex dendritic structure and are more likely to receive higher frequency and larger amplitude excitatory events. As such, it is possible that these events can facilitate the activation of male CART-negative MCH cells.

Overall, this study expanded on the heterogeneity of the MCH system and generated brain atlas templates to guide future work targeting different MCH cells. Neurochemical and electrical distinctions supported the division of MCH/CART+ and MCH/CART– cells as unique subpopulations, and novel transgenic tools to selectively isolate CART-positive and -negative MCH cells will be helpful in elucidating the complex and multifaceted contributions of the MCH system.

## ACKNOWLEDGMENTS

This work is supported by a NSERC Discovery Grant (MJC), I-CUREUS (PAM, JWI), NSERC Canada Graduate Scholarship (ASS), Ontario Graduate Scholarship (BHH). The authors thank Dr. Matthew Holahan for access to the Neurolucida platform.

## TABLES

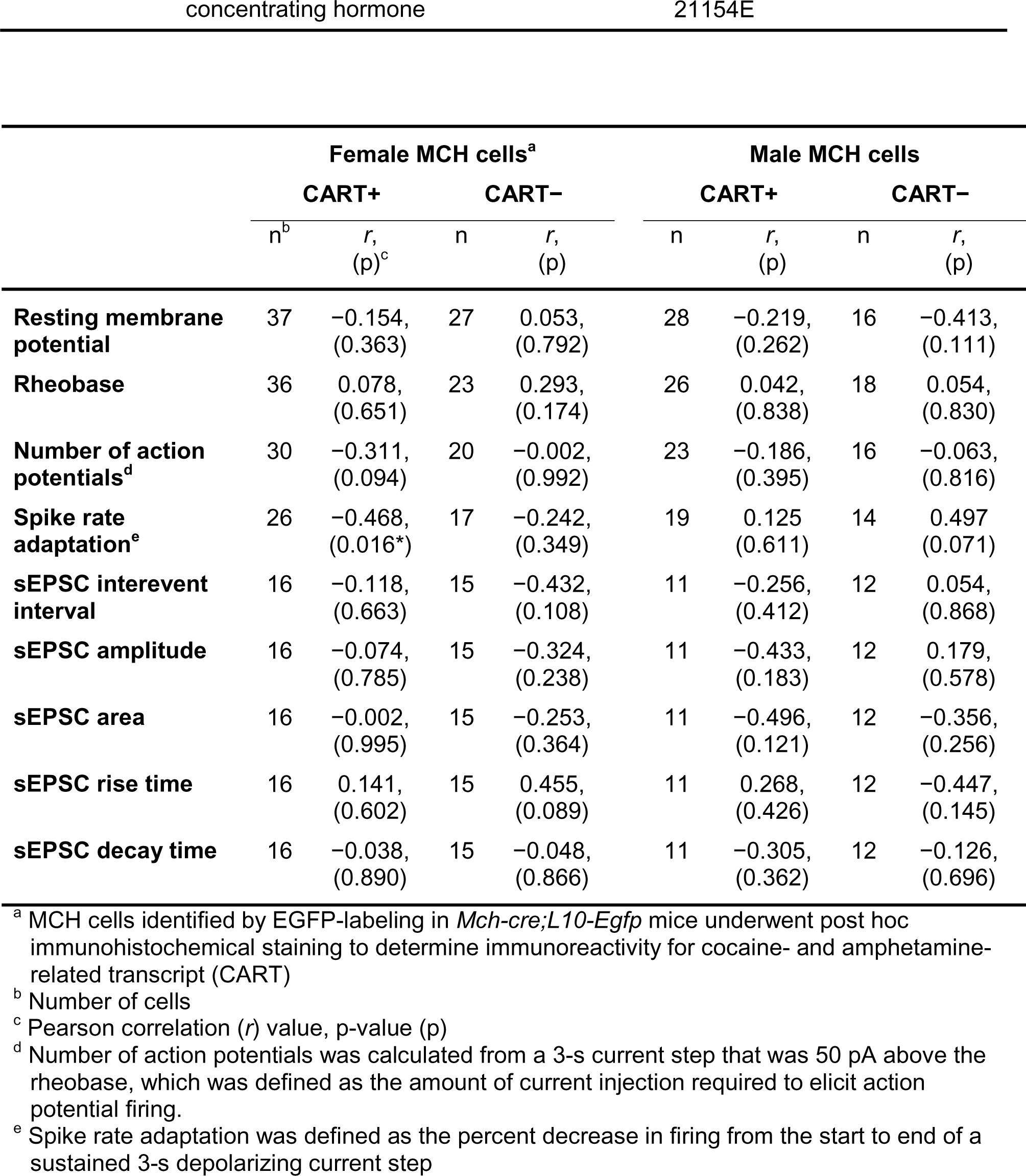
Supplemental Table 1. Electrophysiological features at MCH cells in relation to age of male and female mice.

## Notes

### Competing Interest Statement

The authors have declared no competing interest.

