## Supplemental Table 1 for "Neuroanatomical, electrophysiological, and morphological characterization of melanin-concentrating hormone cells coexpressing cocaine- and amphetamine-regulated transcript"

**Supplemental Table 1. Electrophysiological features at MCH cells in relation to age of male and female mice.**

|  | **Female MCH cells^a^** | | | |  | **Male MCH cells** | | | |
| --- | --- | --- | --- | --- | --- | --- | --- | --- | --- |
|  | **CART+** | | **CART−** | |  | **CART+** | | **CART−** | |
|  | n^b^ | *r*,  (p)^c^ | n | *r*,  (p) |  | n | *r*,  (p) | n | *r*,  (p) |
| **Resting membrane potential** | 37 | −0.154,  (0.363) | 27 | 0.053,  (0.792) |  | 28 | −0.219,  (0.262) | 16 | −0.413,  (0.111) |
| **Rheobase** | 36 | 0.078,  (0.651) | 23 | 0.293,  (0.174) |  | 26 | 0.042,  (0.838) | 18 | 0.054,  (0.830) |
| **Number of action potentials^d^** | 30 | −0.311,  (0.094) | 20 | −0.002,  (0.992) |  | 23 | −0.276,  (0.412) | 16 | 0.054,  (0.868) |
| **Spike rate adaptation^e^** | 26 |  | 18 |  |  | 22 |  | 16 |  |
| **sEPSC interevent interval** | 16 | −0.118,  (0.663) | 15 | −0.432,  (0.108) |  | 11 | −0.186,  (0.395) | 12 | −0.063,  (0.816) |
| **sEPSC amplitude** | 16 | −0.074,  (0.785) | 15 | −0.324,  (0.238) |  | 11 | −0.433,  (0.183) | 12 | 0.179,  (0.578) |
| **sEPSC area** | 16 | −0.002,  (0.995) | 15 | −0.253,  (0.364) |  | 11 | −0.496,  (0.121) | 12 | −0.356,  (0.256) |
| **sEPSC rise time** | 16 | 0.141,  (0.602) | 15 | 0.455,  (0.089) |  | 11 | 0.268,  (0.426) | 12 | −0.447,  (0.145) |
| **sEPSC decay time** | 16 | −0.038,  (0.890) | 15 | −0.048,  (0.866) |  | 11 | −0.305,  (0.362) | 12 | −0.126,  (0.696) |
| ^a^ MCH cells identified by EGFP-labeling in *Mch-cre;L10-Egfp* mice underwent post hoc immunohistochemical staining to determine immunoreactivity for cocaine- and amphetamine-related transcript (CART)  ^b^ Number of cells  ^c^ Pearson correlation (*r*) value, p-value (p)  ^d^ Number of action potentials was calculated from a 3-s current step that was 50 pA above the rheobase, which was defined as the amount of current injection required to elicit action potential firing.  ^e^ Spike rate adaptation was defined as the percent decrease in firing from the start to end of a sustained 3-s depolarizing current step | | | | | | | | | |
