## Supplemental Figure 1 for "Neuroanatomical, electrophysiological, and morphological characterization of melanin-concentrating hormone cells coexpressing cocaine- and amphetamine-regulated transcript"

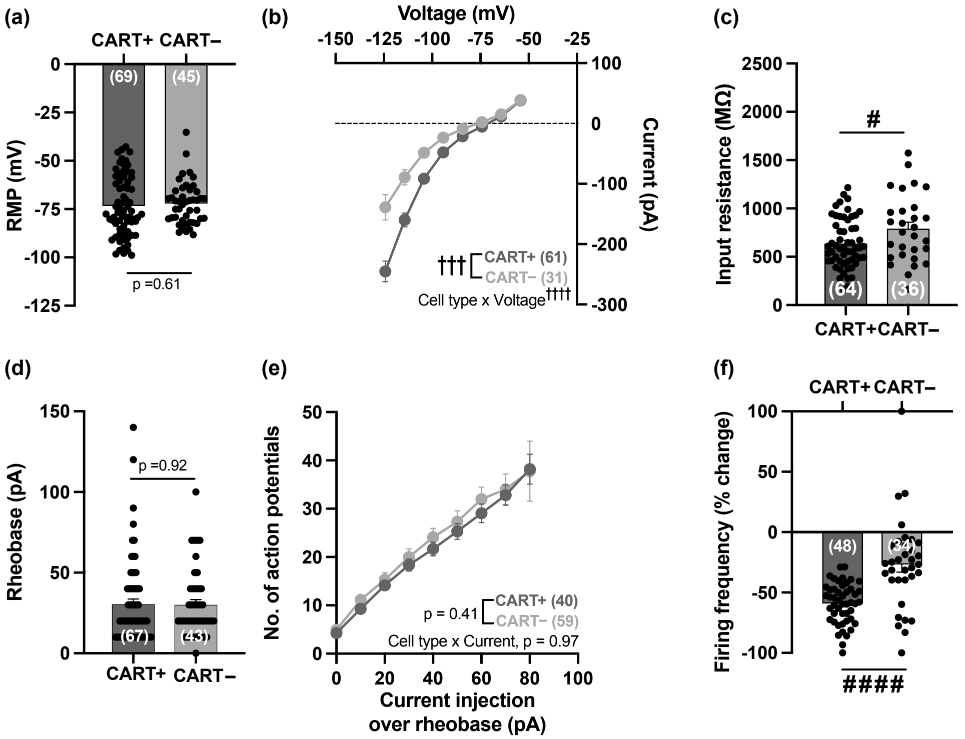


**Supplemental Figure 1. MCH/CART+ cells exhibited lower input resistance and greater spike rate adaptation.** Comparison of resting membrane potential (RMP; ***a***), current-voltage relation (***b***), input resistance (***c***), rheobase (***d***), number of action potentials elicited (***e***), and percent change in firing frequency of MCH/CART+ and MCH/CART−. Two-way ANOVA: †††, $p < 0.001$; ††††, $p < 0.0001$. Unpaired t test: #, $p < 0.05$; # # # #, $p < 0.0001$.
