## Supplemental Figure 2 for "Neuroanatomical, electrophysiological, and morphological characterization of melanin-concentrating hormone cells coexpressing cocaine- and amphetamine-regulated transcript"

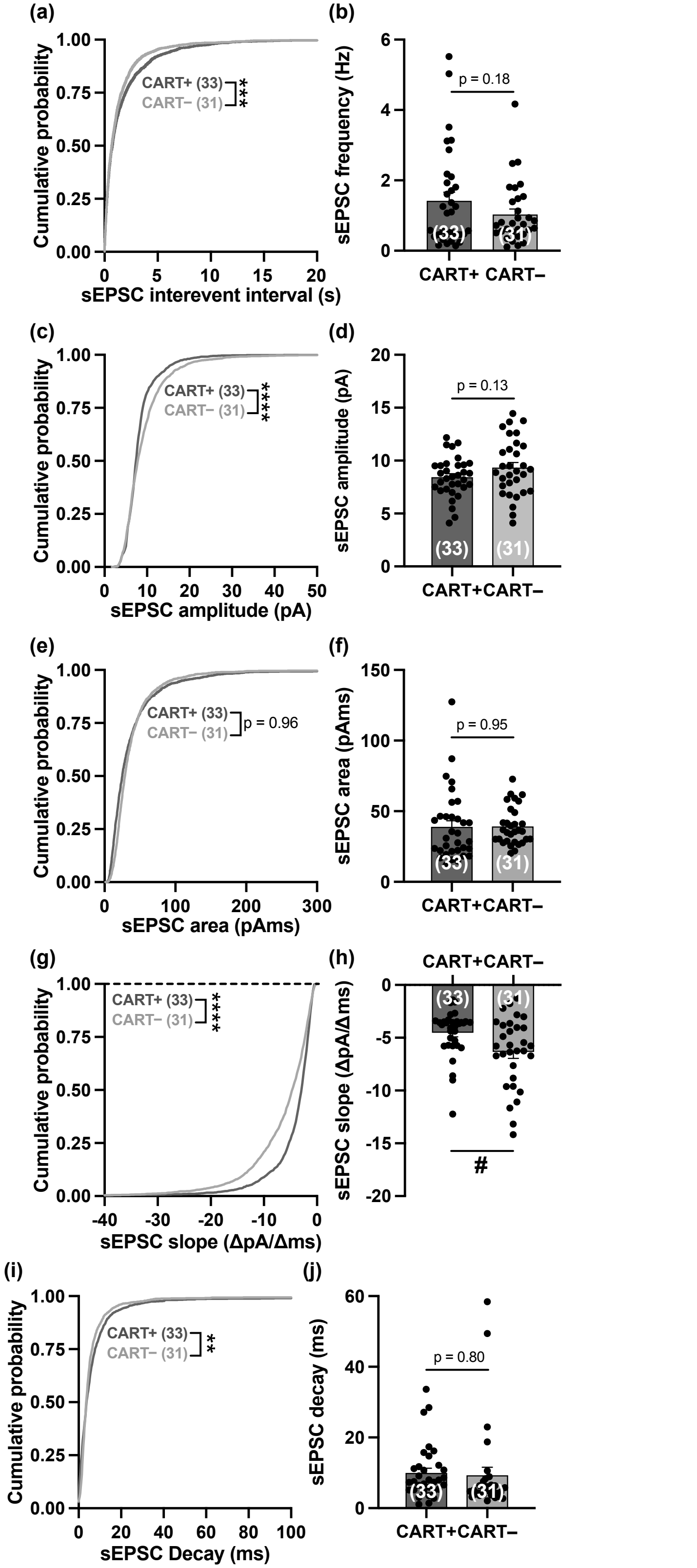


**Supplemental Figure 2. MCH/CART+ cells receive smaller and less frequent sEPSC events.** Cumulative probability plots and average values of sEPSC event interevent interval and frequency (***a***, ***b***), amplitude (***c***, ***d***), area (***e***, ***f***), slope (***g***, ***h***), and decay (***i***, ***j***) recorded from MCH/CART+ and MCH/CART− cells. Unpaired t test: *, $p < 0.05$. Kolmogorov-Smirnov test: **, $p < 0.01$; ***, $p < 0.001$; ****, $p < 0.0001$.
