## Supplemental Figure 3 for "Neuroanatomical, electrophysiological, and morphological characterization of melanin-concentrating hormone cells coexpressing cocaine- and amphetamine-regulated transcript"

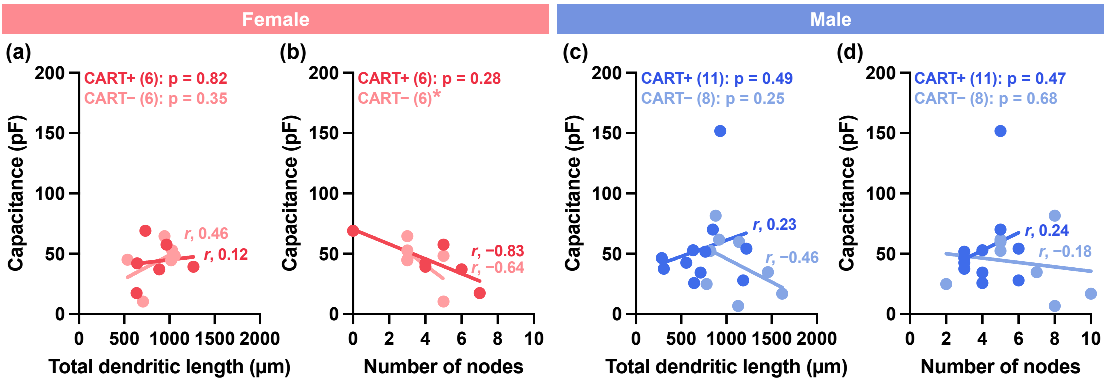


**Supplemental Figure 3. Calculated membrane capacitance was unrelated to the dendritic structure of MCH cells.** Correlation plot and Pearson correlation value (*r*) of membrane capacitance as a function of total dendritic length (***a***, ***c***) and the number of nodes (***b***, ***d***) at female and male MCH/CART+ and MCH/CART– cells.
