## Supplemental Figure 4 for "Neuroanatomical, electrophysiological, and morphological characterization of melanin-concentrating hormone cells coexpressing cocaine- and amphetamine-regulated transcript"

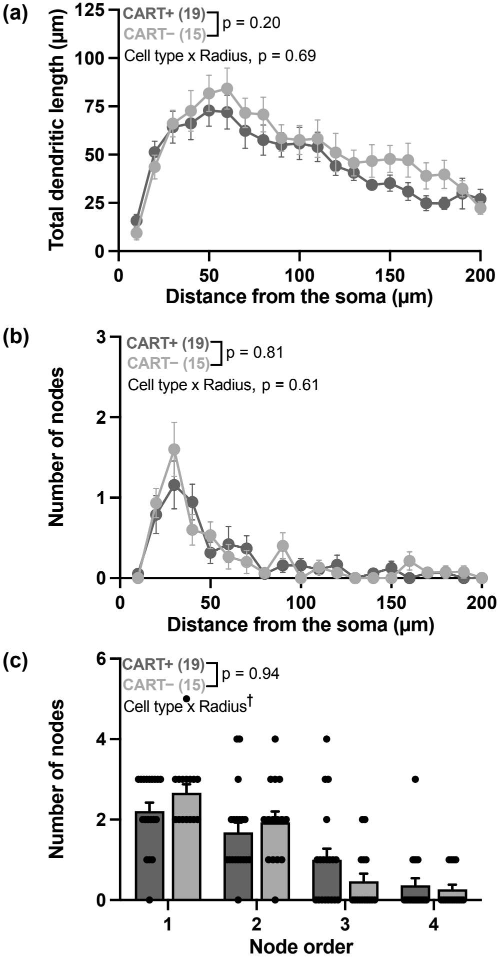


**Supplemental Figure 4. No morphological differences between MCH/-CART+ and MCH/CART– cells.** Comparison of total dendritic length (***a***), the number of nodes at each node order (***b***), and the number of nodes (***c***) measured within a 200 mm radius from the soma of the MCH/CART+ and MCHR/CART– cells. Two-way ANOVA: †, $p < 0.05$.
